# Towards Clinical Phage Microbiology for Biofilm Infections: A Comparative Study and a Suggested Protocol

**DOI:** 10.1101/2025.03.07.642007

**Authors:** Amit Rimon, Ron Braunstein, Ortal Yerushalmi, Noa Katz, Lidor Yosef, Yitzchak Gvili, Hadil Onalla, Shunit Coppenhagen-Glazer, Ran Nir Paz, Ronen Hazan

## Abstract

Non-resolving bacterial infections involve biofilm formation and are often complicating treatments. Utilizing lytic bacteriophages with antibiotics holds promise in biofilm eradication. Accurately matching phage-antibiotic combinations against target bacteria, termed Clinical Phage Microbiology (CPM), is crucial in effective phage therapy treatments. However, compared to planktonic cultures, performing CPM on biofilm infections poses a significant challenge due to the lack of effective methods and a standard protocol. This study compared various CPM approaches in biofilms. To this end, the activity of five phages was assessed against *Pseudomonas aeruginosa* biofilms using nine methods and multiple approaches. Here, we discuss various aspects of each technique, including sensitivity, duration, ease of implementation in diagnostic labs, and labor. Finally, we offer a preliminary protocol for testing phage-sensitivity in biofilm, which was tested on various bacteria species, and may serve as a basis for comprehensive CPM in biofilm.

## Introduction

The escalating threat of antibiotic-resistant bacterial infections constitutes a critical challenge to global health. It is ranked as the second leading cause of mortality today and is anticipated to become the primary cause of death by 2050(Ikuta et al., 2022; Murray et al., 2022). One significant contributor to these persistent infections is the formation of bacterial biofilms, which are structures composed of microorganisms embedded in an extracellular matrix. Unlike individual bacterial cells, these biofilms exhibit remarkable resilience to conventional antibiotics due to antibiotic-limited penetration capabilities(Ciofu et al., 2017; Parrish & Fahrenfeld, 2019), making these infections particularly challenging to treat(Stewart & Costerton, 2001). Some examples of such clinical conditions are osteomyelitis(Lew & Waldvogel, 2004; Yun et al., 2008), diabetic foot ulcers(Pouget et al., 2020), and medical device-related infections, including catheter-associated UTIs(Nickel & Costerton, 1992),(Akers et al., 2014; Yun et al., 2008) Moreover, biofilms pose a considerable threat in many fields, including the food industry(Galie et al., 2018), agriculture(R. Bhatia et al., 2021), water systems(S. Liu et al., 2016) and many more.

A promising avenue in the battle against biofilms is the utilization of bacteriophages (phages), viruses that invade bacterial cells, exploiting their systems to propagate, and ultimately kill the bacterial lysis followed by spreading of hundreds of new phage particles, sicking for more cells. Phage therapy demonstrated efficacy in compassionate treatments(McCallin et al., 2019; Nir-Paz et al., 2019; Onallah et al., 2023; Yerushalmy et al., 2023) in *in vivo* studies(Pires et al., 2022; Rimon et al., 2023) and offers several advantages over antibiotics. Phages are highly strain-specific with minimal impact on the host’s commensal microbiome. They also exhibit “Auto-Dosing” kinetics correlated with their target, and are relatively easy to isolate and genetically improve. Additionally, certain phages carry depolymerase, enzymes capable of degrading the extracellular matrix, enhancing their ability to penetrate and dismantle biofilm layers(Azeredo & Sutherland, 2008; Loc-Carrillo & Abedon, 2011; Pires et al., 2022; Rimon et al., 2023).

However, a significant hurdle in phage therapy lies in the precise matching of suitable phages to the patient’s bacterium. Various phages, alone or in combination with antibiotics, are evaluated through Clinical Phage Microbiology (CPM) methods on planktonic bacteria(Gelman et al., 2021; Yerushalmy et al., 2023). However, a validated and reliable method for assessing the impact of different phages on biofilms is currently lacking(Abedon et al., 2021; Ferriol-González & Domingo-Calap, 2020; Pires et al., 2022) Consequently, the phage-matching process for biofilm-associated infections relies on testing phages’ efficacy against bacteria in their planktonic state or utilizing plaque plate assays.

In this study, we addressed this critical gap by comparing nine methods that have demonstrated potential for quantifying the growth of biofilms in non-phage contexts(Kırmusaoğlu, 2019; Peeters et al., 2008), phage activity in biofilms. To this end, we used five *Pseudomonas aeruginosa* (*P. aeruginosa*) phages(Rimon et al., 2025). In planktonic and biofilm cultures. These methods included live-dead staining, Crystal Violet staining, and Colony Forming Units (CFU) assay following sonication of biofilms(Kragh et al., 2019), calorimetry(Lichtenberg et al., 2022; Wadsö et al., 2017), oxygen flux measurement(S. Bhatia et al., 2021), qPCR for extracellular DNA(Hazan et al., 2016a), extracellular ATP measurement(Ihssen et al., 2021) and biomass assessment using fluorescent dye and Scanning Electron Microscopy (SEM). Finally, we describe a possible protocol, which was assessed on different bacterial species: another *P. aeruginosa* strain, *Escherichia coli* (*E. coli*), *Staphylococcus aureus* (*S. aureus*), and *Enterococcus faecalis* (*E. faecalis*).

## Results

### A. The problem: Growth curve analysis is not fit for biofilm analysis

The most frequently employed method for evaluating bacterial growth inhibition by phages is through growth curve analysis(Yerushalmy et al., 2023). We have assessed all of our phages in two different titers on planktonic bacteria, to ensure variable potency. The area under the curve (AUC) varies significantly among different phages at similar MOI levels (*p*-value<0.05, Fig. 1a-f). Additionally, four out of five phages, except PADD, exhibit increased activity at higher MOI levels (*p*-value<0.05, Fig. 1a-f). However, it is not as efficient when assessing biofilm bacteria, due to the visibility obstruction caused by adhered bacteria(Yerushalmy et al., 2023) Nevertheless, we have undertaken this assessment in this study. Optical density (OD_600_ nm) measurements varied considerably across the majority of the phages tested, with only PASA16 and PADD demonstrating discernible differences among the five phages evaluated (Fig. 1g-h). This indicates the necessity to identify alternative methods for monitoring phage efficacy in biofilm-grown bacteria.

**Figure 1:**
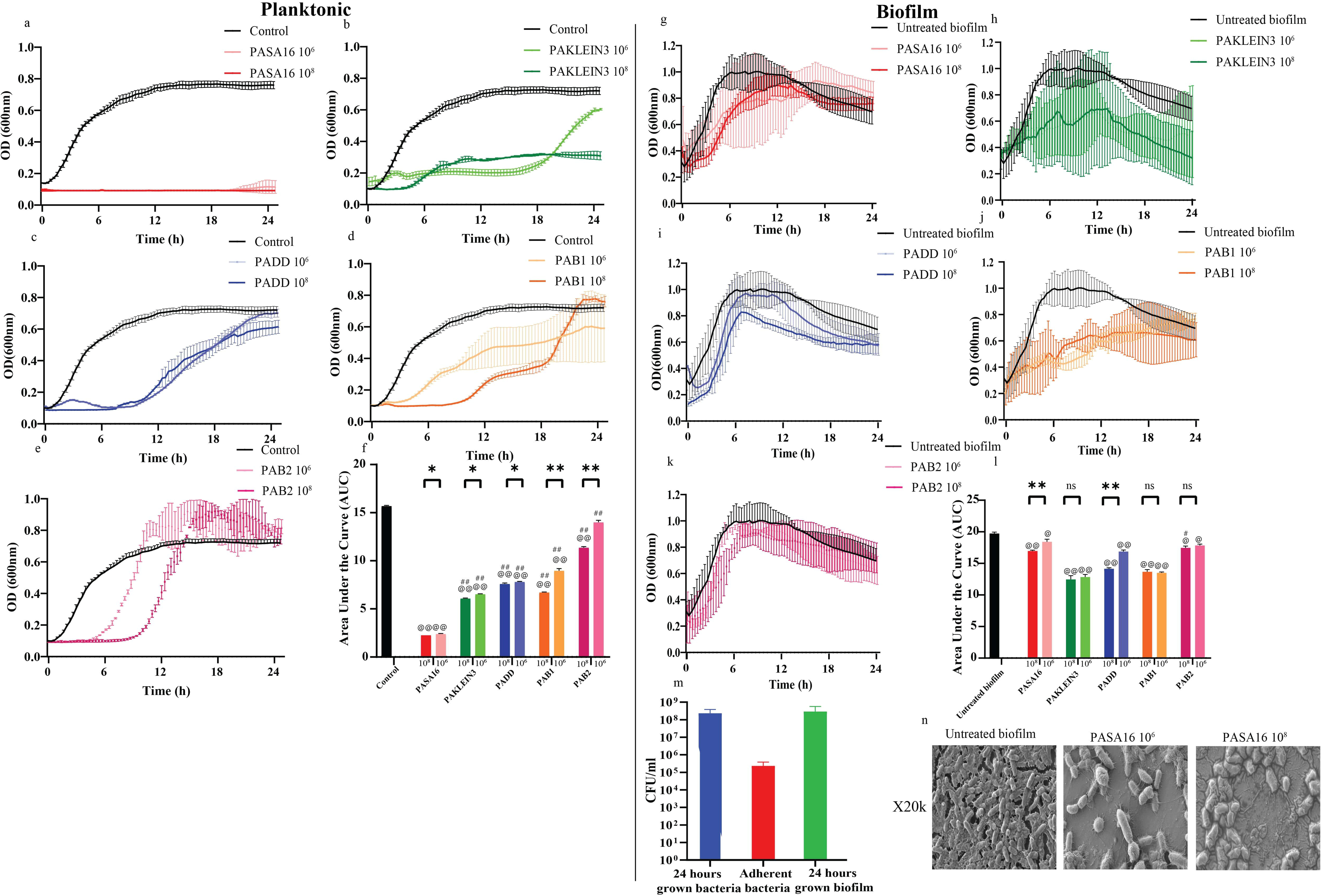
Growth curves of all phages on planktonic and biofilm bacteria and model establishment. Optical density (OD_600_ nm) of *P. aeruginosa* bacteria in **a-f** planktonic cultures or **g-l** biofilm treated with five different phages at PFU concentrations of 10^8^ and 10^6^. Phages include **a, g** PASA16 phage, **b, h** PAKLEIN3 phage, **c, i** PADD phage, **d, j** PAB1 phage, **e, k** PAB2 phage. OD was measured every 20 minutes for 24 hours. **f, l** The area under the curve (AUC) of each growth curve was calculated at different concentrations. **m** To validate true biofilm formation, CFU counts were compared between 24-hour-grown planktonic bacteria, 24-hour-grown biofilm (washed), and 10-second bacterial adhesion (washed similarly to pre-grown biofilm). Higher CFU counts in pre-grown biofilms confirmed true biofilm formation rather than mere bacterial adherence. **n** Scanning Electron Microscopy (SEM) was done to assess biofilm formation in our model and to support the assumption that PASA16 was indeed the most effective phage (all phages are shown in Supplemental Figure S5). The results are the average of triplicates, presented as mean ± standard deviation. Representative results from two independent experiments. * *p <* 0.05, ** *p <* 0.001 by two-way Student’s t-test, comparing different titers of a given phage. # *p <* 0.05, ## *p <* 0.001 by one-way Student’s t-test, evaluating whether a given phage is significantly less effective than PASA16. @ *p <* 0.05, @@ *p <* 0.001 by one-way Student’s t-test, assessing whether each phage is significantly more effective than the control.

### B. Experimental design and biofilm model establishment

*Pseudomonas aeruginosa* biofilms were cultivated in 96-well plates using a previously established protocol. To ensure the integrity of the biofilm and remove any loosely adherent bacteria, the wells were gently washed with fresh media before phage treatment. To confirm that the bacterial growth represented true biofilm formation rather than merely adhered planktonic cells, we conducted a verification process. In parallel to biofilm cultivation, freshly grown overnight planktonic bacterial cultures were added to separate wells. Upon enumerating the colony-forming units (CFU), we found that the planktonic control had a CFU count approximately three orders of magnitude lower than that of the pre-grown biofilm. This significant difference in bacterial density conclusively demonstrated that the observed growth in the biofilm-cultivated wells represented a true, mature biofilm structure, thereby ensuring the reliability and relevance of subsequent phage treatment experiments (Fig. 1m) also supported by the SEM results (Fig. 1n).

To evaluate phage activity on *Pseudomonas aeruginosa* biofilm, we utilized five *Pseudomonas aeruginosa*-targeting phages from the Israeli Phage Bank (Yerushalmy et al., 2020). These phages, which are detailed in other publications(Rimon et al., 2025) include PASA16, PAKlein3, PADD, PAB1, and PAB2 (Table 1). PASA16, in particular, is renowned for its involvement in numerous phage therapy compassionate cases worldwide (Alkalay-Oren et al., 2022; Khatami et al., 2021; Onallah et al., 2023).

**Table 1.**
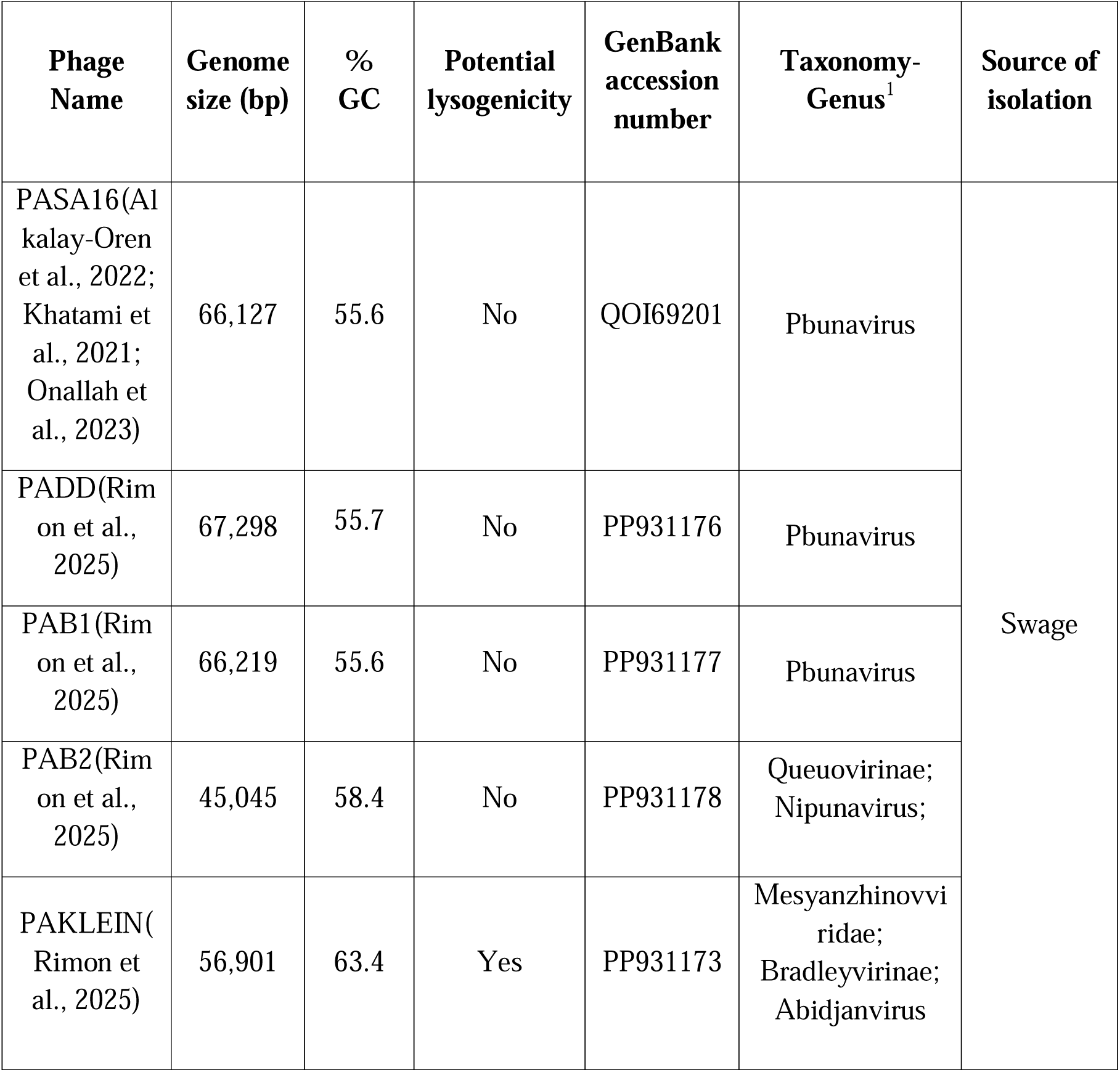
Description of the *pseudomonas aeruginosa* targeting phages used in this study. The phages used here are from the Israeli Phage Bank (IPB(Yerushalmy et al., 2020)), and their isolation location and genomic properties were described(Rimon et al., 2025). This table details various genomic characteristics of different phages, including the genome length, GC content (%), potential for lysogenicity determined by the presence of repressor proteins, and the presence of phage genomes in bacterial genomes identified through BLAST comparisons. Potential for temperate life cycle determined by PhaBOX. ^1^ *Viruses; Duplodnaviria; Heunggongvirae; Uroviricota; Caudoviricetes*

We hypothesized that a method sensitive enough to differentiate between various conditions would also accurately distinguish between different phages. To test this, we evaluated each phage at two different titers, as it is generally accepted that a phage used at a higher titer is more effective than the same phage at a lower titer (Meneses et al., 2023).

Our experimental design included comparing each phage-treated group to an untreated control group. Given PASA16’s established efficacy in clinical settings and our preliminary SEM analysis, we sought a method that would demonstrate its superior performance (Fig. 1n).

### B. Common methods: Sonication-CFU, Crystal Violet, Live Dead stains

First, we examined methods that are currently in use for biofilm assessment, such as Colony Forming Unit (CFU), Crystal Violet (CV), and Live dead stain(Kragh et al., 2019).

#### 1. CFU assay

CFU is conducted post-sonication of the biofilm, and measuring bacterial cell viability. However when using phages at a concentration of 10^8^ PFU/ml, a significant reduction in comparison to the untreated biofilm control was observed in 70% of phages tested (*p-value* <0.05, Fig. 2a). Notably, the CFU assay successfully differentiated between different phage concentrations for 60% of the tested phages, and in comparison to PASA16 only 25% of phages were inferior (*p-value*<0.05, see Fig. 2a). Interestingly, when CFU measurements were modified and taken from an stainless steel washer rather than directly from the 96-well plate, the assay showed improved ability at all three parameters described (Fig. 2b).

**Figure 2.**
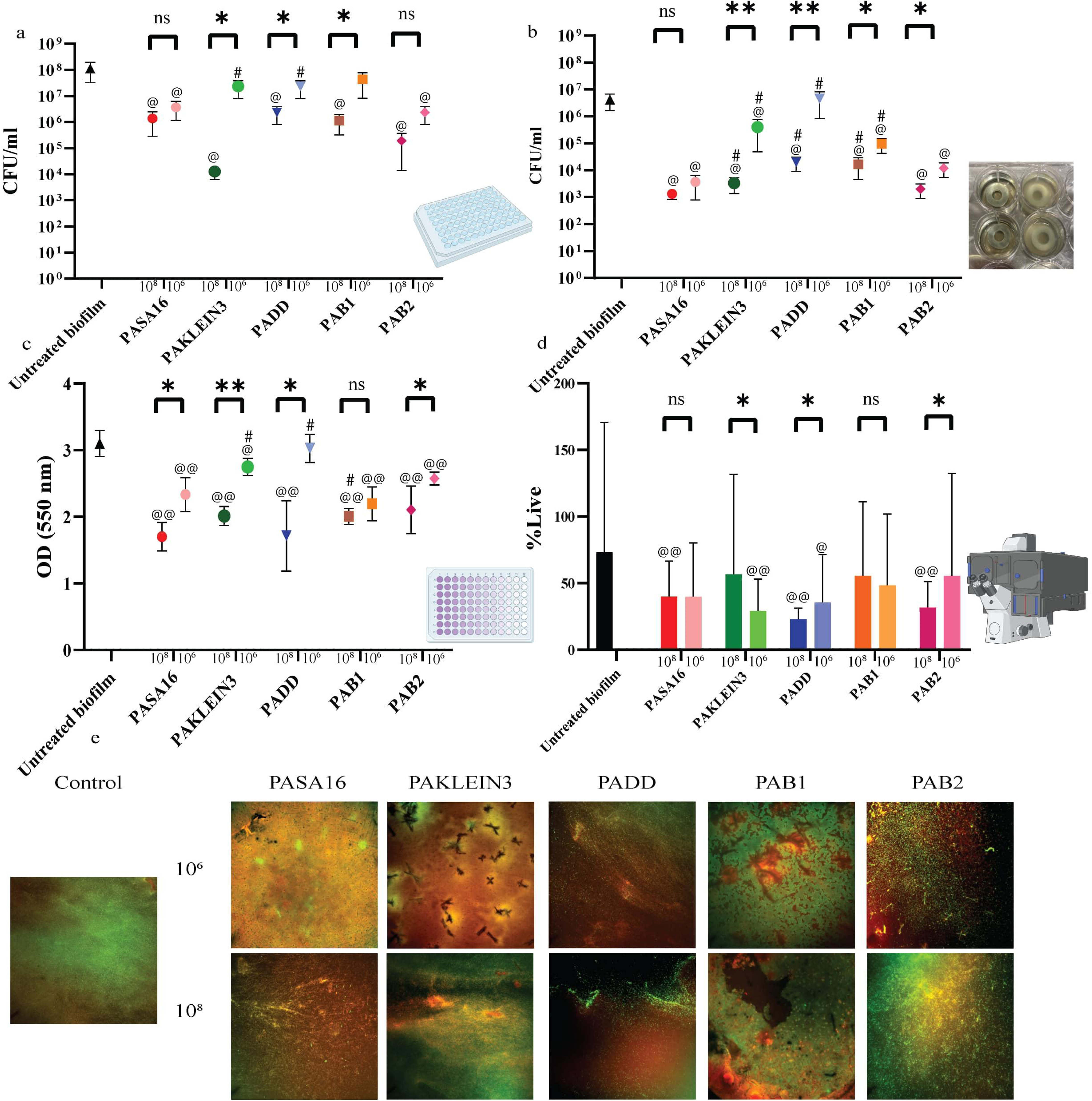
Assessment of pre-grown biofilm using CFU, crystal violet, and live-dead analysis. Biofilm subjected to treatment with five different phages at two concentrations (10□ and 10□ PFU/ml) was evaluated through multiple assays: **a** Quantification of colony-forming units (CFU) after sonication. **b** Evaluation of CFU after sonication with the use of washers. **c** utilization of crystal violet staining for biofilm assessment. **d** Quantitative evaluation of Live staining (GFP) with the calculation of average % live bacteria of the total, following the methodology by Robertson et al(Robertson et al., 2019). **e** Live-dead staining analysis utilizing Live (CYTO9) in green and Dead (Propidium iodide) in red, captured at 10X magnification. The results are the average of triplicates, presented as mean ± standard deviation. Representative results from two independent experiments. * *p <* 0.05, ** *p <* 0.001 by two-way Student’s t-test, comparing different titers of a given phage. # *p <* 0.05, ## *p <* 0.001 by one-way Student’s t-test, evaluating whether a given phage is significantly less effective than PASA16. @ *p <* 0.05, @@ *p <* 0.001 by one-way Student’s t-test, assessing whether each phage is significantly more effective than the control. This figure was made using biorender.com

#### 2. The Crystal Violet (CV) assay

This assay focuses on biomass rather than living cells, performed at experiment termination as an endpoint analysis, and demonstrated sensitivity in discerning between the activities of different concentrations for 80% of phages tested (*p-value* <0.05, Fig. 2c). When comparing different phages with similar titers, only one phage (PAB1) is not significantly different from the same phage a lesser titer (90% are superior to untreated control and 37.5% are inferior to PASA16 (p-value < 0.05, Figure 2c).

#### 3. Live Dead stain

This dye is used for visualizing bacterial viability. First assessed for phage and bacterial toxicity. Although SYTO9 presented mild cytotoxicity, it was not phage-toxic (*p-value*<0.05, non-significant accordingly, Supplemental Fig. S1a-b). Consequently, all analyses were conducted at the endpoint and not read online. The percentage of live bacteria within the total population indicated a significant reduction for 50% of phages (*p-value*<0.05 Fig. 2d, e). Remarkably, only 40% out of the five tested phages exhibited a correlation where higher phage titers resulted in lower %live, and 0% were superior to PASA16. (*p-value*<0.05, Fig. 2d, e).

It is noteworthy that all these methods require the termination of the experiment and cannot be assessed in real time. However, modified CFU measurements using washers remain a method sensitive enough for future phage differentiation. Alongside Crystal Violet is sensitive when comparing different titers, and when comparing phages to control, but its ability to detect inter-phage variability is limited. These two methods complement each other, with one assessing bacterial viability and the other evaluating biomass. Importantly, both techniques are easy to implement and do not require specialized equipment or materials (Table 2).

**Table 2.**
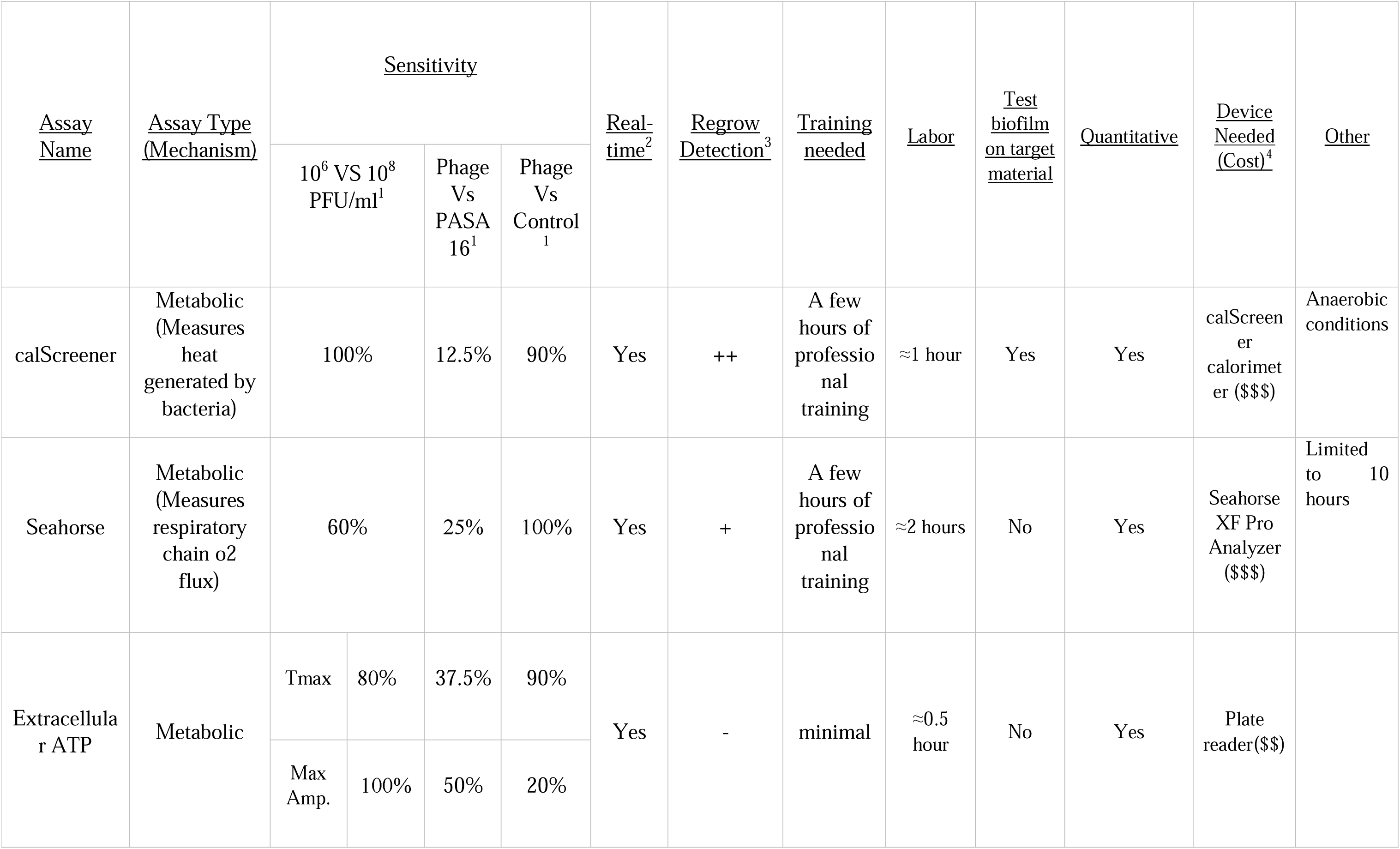
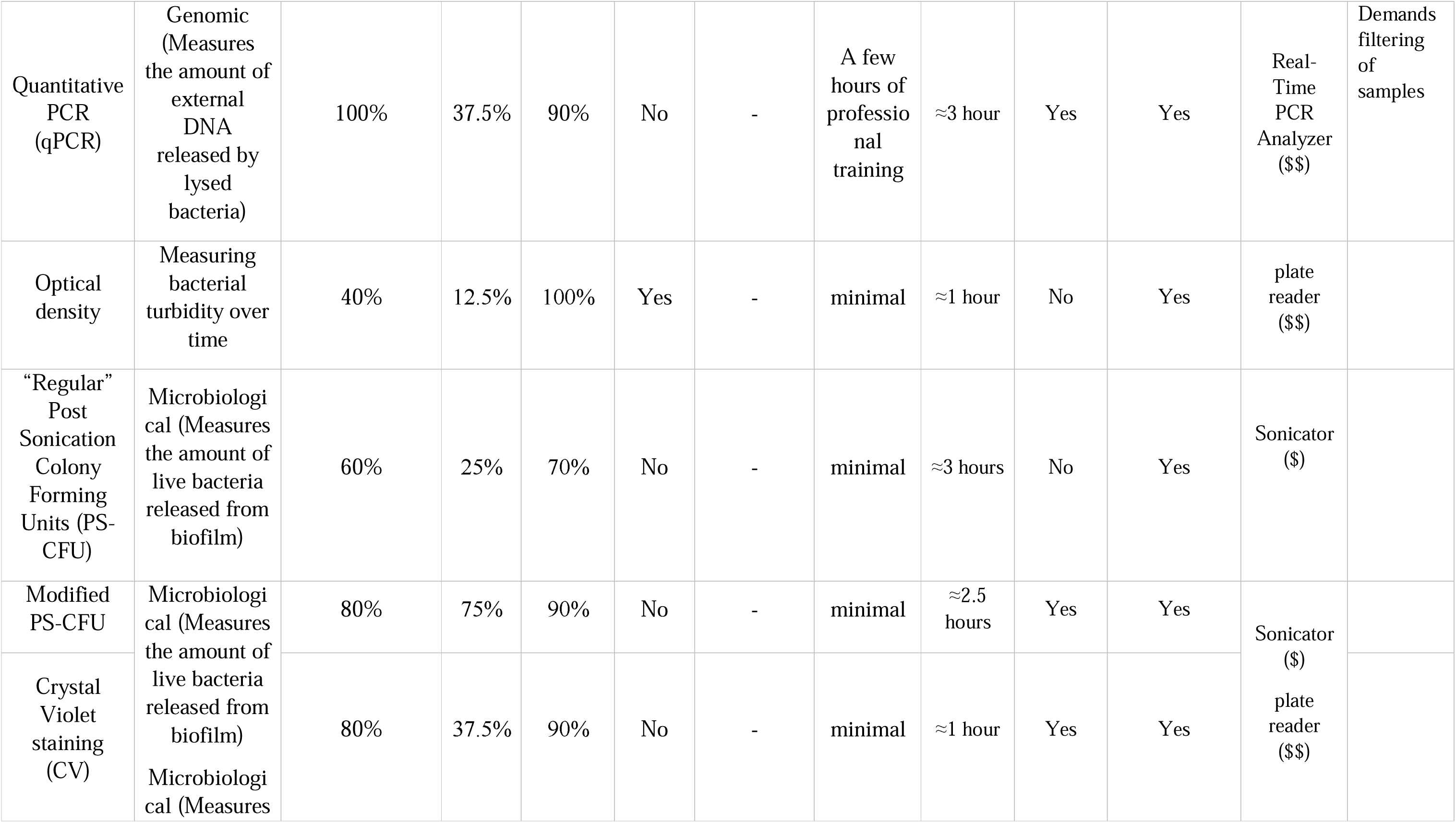
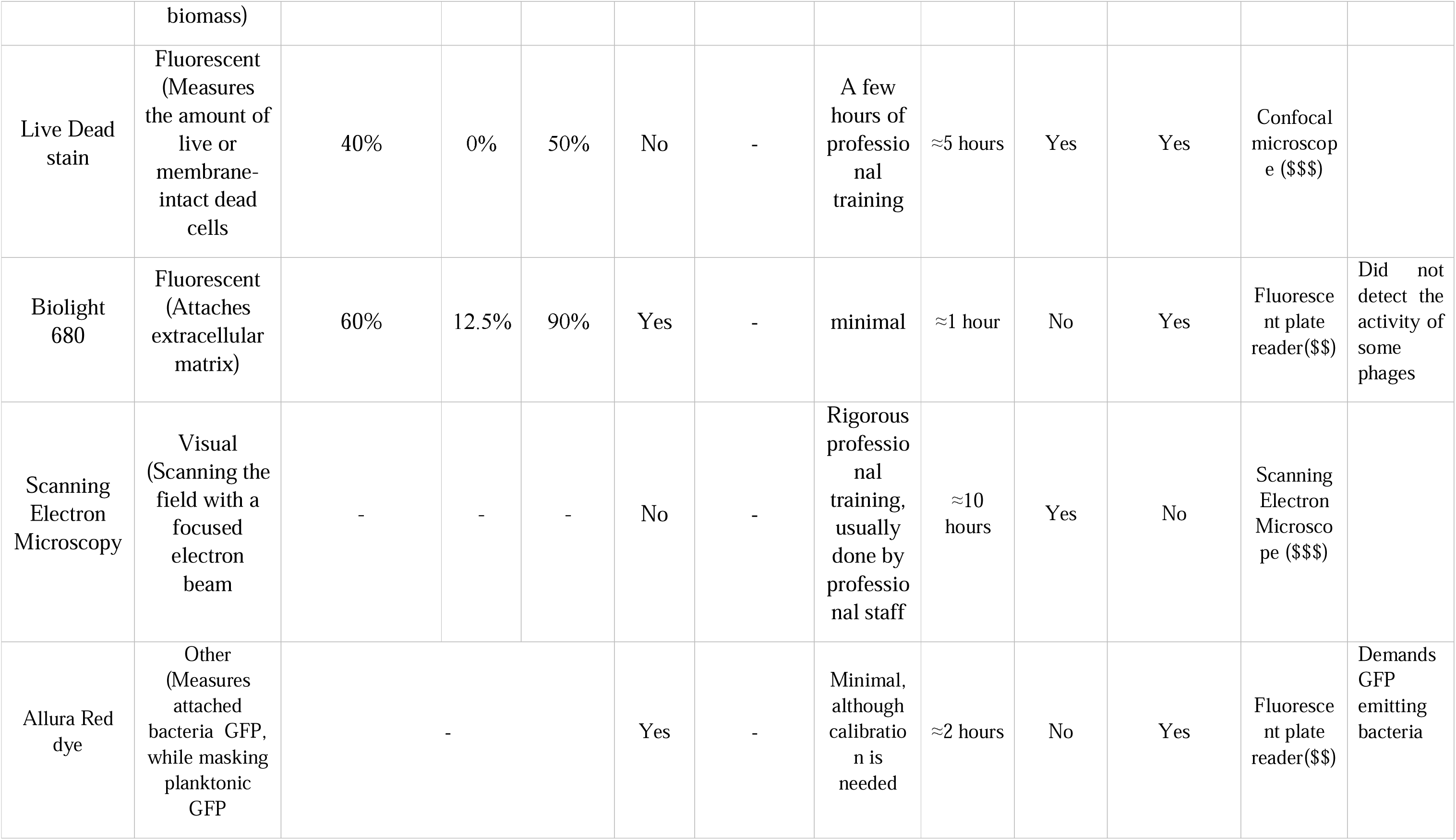
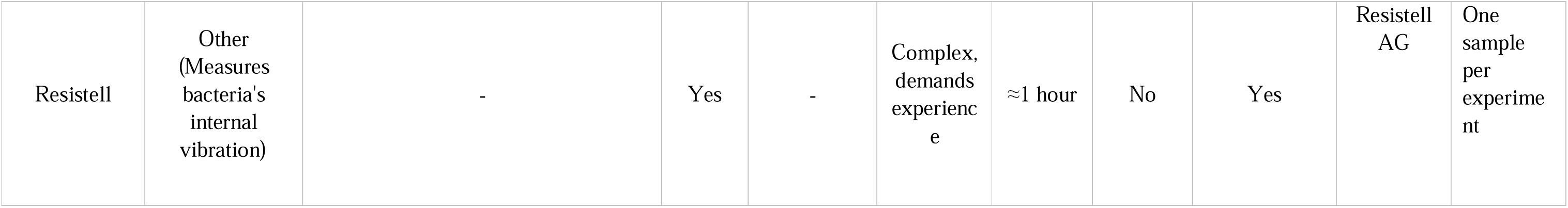
Summary of various characteristics of the evaluated phage efficacy on biofilm methods. Each assay is described in terms of its name, type, mechanism, and sensitivity (ranked between 1-5)^1^. The table also assesses the ability of each test to provide real-time data^2^, detect bacterial regrowth^3,^ test on target material, and quantify results. Additionally, the table includes rankings for training requirements and associated costs^4^ for each method. PS Post-Sonication, Tmax “Time to maximal amplitude”, Max Amp “Maximal Amplitude” ^1^10^6^ VS 10^8^ PFU/ml -The % of phages, out of 5 phages tests, in which the method differentiates between different titers. Phage Vs PASA16 – the % of phages that are significantly inferior to PASA16, which is established as the most efficient phage, out of 4 phages. Phage Vs Control – The % of phages that significantly reduce bacterial count or activity, in comparison to the untreated biofilm, out of 5 phages. ^2^Real time – a method that enables bacterial monitoring live through the experiment, and not only at the endpoint. ^3^The ability to detect bacterial regrowth after initial reduction. ^4^$ - basic lab equipment; $$ - specialized lab equipment; $$$ - unique machinery commonly obtained only by core facilities.

### C. Metabolic assessments

In contrast to optical methods, metabolic assays do not require a complete well-read-through. They use other quantification parameters such as temperature changes, pH changes, and ATP. We first aimed to quantify bacteria using the metabolic activity of living cells. Which can be obstructed by aggregated bacteria. Three different methods were used, calorimetry, PH measurement, and extracellular ATP.

#### 4. Calorimetry assessment

Enables the quantification of heat produced, typically associated with metabolic changes in cell cultures(Lichtenberg et al., 2022; Wadsö et al., 2017) In this study, we utilized the CalScreener® (https://symcel.com/) calorimeter, which operates within a closed 32-well plate system, to measure heat flow in each well. A notable difference between control and phage treatment was primarily observed within the initial 10 hours of measurement (Fig. 3a-e). Introducing phages at 10^8^ PFU/ml resulted in a noteworthy reduction in AUC, with a 28.8% decrease for PASA16-treated and a 92% reduction for PAB2-treated biofilms compared to the untreated control (*p-value* <0.001 using 2-way Student t-test, Fig. 3a-e). Intriguingly, the AUC for all 100% of phages was higher when treated with lower phage PFU compared to higher phage titers, and 90% of phages were superior to untreated control. On the other hand, only 12.5% of phages were inferior to PASA16 (*p-value<*0.001, Fig. 3a-f). A decline in metabolic activity was observed in all CalScreener wells due to the anaerobic growth conditions created by limited oxygen availability. Notably, bacterial regrowth and adaptation were identified in specific cases (Fig. 3 a, b, d). The initiation of heat measurement after 1 hour of inoculation was a result of the CalScreener protocol, necessitating the assembly of post-treatment plate wells into titanium wells, with their lids sealed.

**Figure 3.**
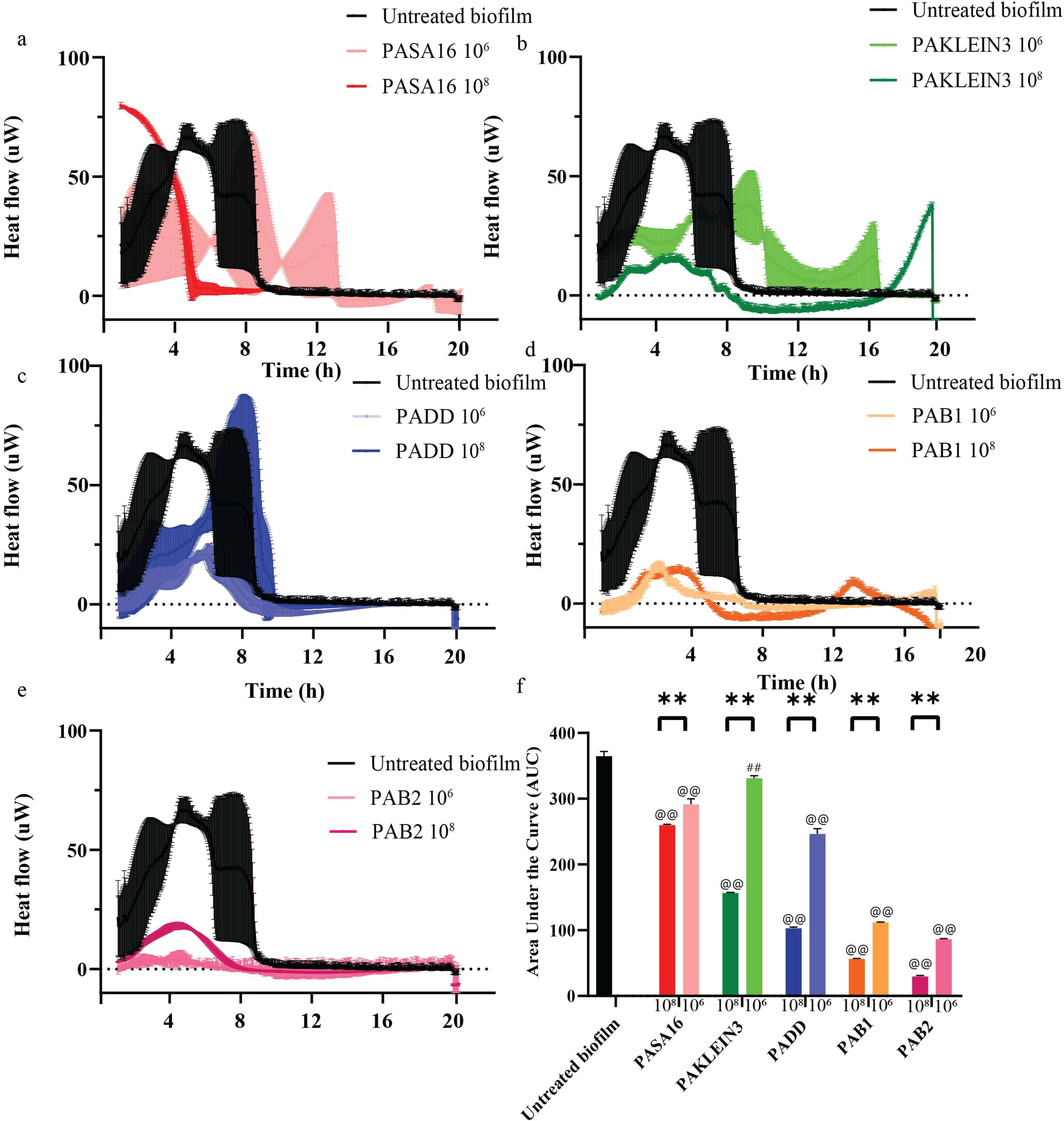
Calorimetric Heat flow-based phage biofilm effect comparison. **a-e** Comparing *P. aeruginosa* biofilm heat generation treated with five different phages at 10^8^ and 10^6^ PFU/ml. **a** PASA16 phage, **b** PAKLEIN3 phage, **c** PADD phage, **d** PAB1 phage, **e** PAB2 phage. **f** The AUC of phage activity was calculated at different concentrations as described above. Notably, a consistent pattern of lower metabolic activity was observed across all five phages when a higher phage titer was applied, a lower AUC was evident when a higher phage titer was utilized by calScreener® calorimetry. The measurement protocol starts 1 hour after inoculation due to the device instructions. The results are the average of triplicates, presented as mean ± standard deviation. Representative results from two independent experiments. * *p <* 0.05, ** *p <* 0.001 by two-way Student’s t-test, comparing different titers of a given phage. # *p <* 0.05, ## *p <* 0.001 by one-way Student’s t-test, evaluating whether a given phage is significantly less effective than PASA16. @ *p <* 0.05, @@ *p <* 0.001 by one-way Student’s t-test, assessing whether each phage is significantly more effective than the control.

#### 5. Oxygen consumption rate (OCR) measurement

This approach measures the amount of oxygen that cells consume over time. The extracellular acidification rate (ECAR) measures the change in pH of the medium surrounding cells over time, reflecting the rate at which cells produce and release acids into the extracellular environment. Both OCR and ECAR can be indicators used to assess cellular metabolism in a biofilm. We employed the Seahorse XFe96 technology, which measures both in a designated microplate(S. Bhatia et al., 2021) Biofilm growth was initiated in the Seahorse XFe96 Analyzer cell plate, and alterations in ECAR were monitored. The AUC of 100% of phage-treated groups exhibited significant differences from that of the untreated control biofilm (*p-value<*0.001 using a 2-way Student t-test, Supplemental Fig. S2a-f). Additionally, in 40% of cases, the AUC decreased among their increased individual tested titers and 25% were inferior to PASA16 (*p-value<*0.001 using 2-way Student t-test, Supplemental Fig. S2f).

#### 6. Extracellular ATP release as an indicator of bacterial lysis

ATP, an essential part of bacterial metabolism, is an intracellular molecule. Therefore, measuring extracellular ATP (eATP) represents both the amount of secreted ATP, as well as ATP spilled from lysed bacteria(Y. Liu et al., 2016). Indeed Carbenicillin lysed PA14 demonstrated a dose-response to eATP levels (Supplemental Fig. S3a).

Measuring eATP, 90% of phages demonstrated a significant reduction in the time needed to reach maximal external ATP levels compared to the untreated control with PASA16 and PAKlein3 showing the most pronounced effect, with 37.5% of phages inferior to PASA16. Furthermore, 80% of phages exhibited higher eATP levels when applied at higher titers. Interestingly, while no phage outperformed PASA16 at 10^8^ PFU/ml, several phages at 10^6^ PFU/ml showed superior performance to this PASA16 concentration (Fig.4 a-f, *p-value<*0.05, using 2-way Student t-test). When measuring the maximal luminescence, 100% of phages showed a dose-response, and 50% of them were inferior to PASA16, but only 25% were superior to the untreated control (Fig.4 a-g, *p-value<*0.05).

**Figure 4.**
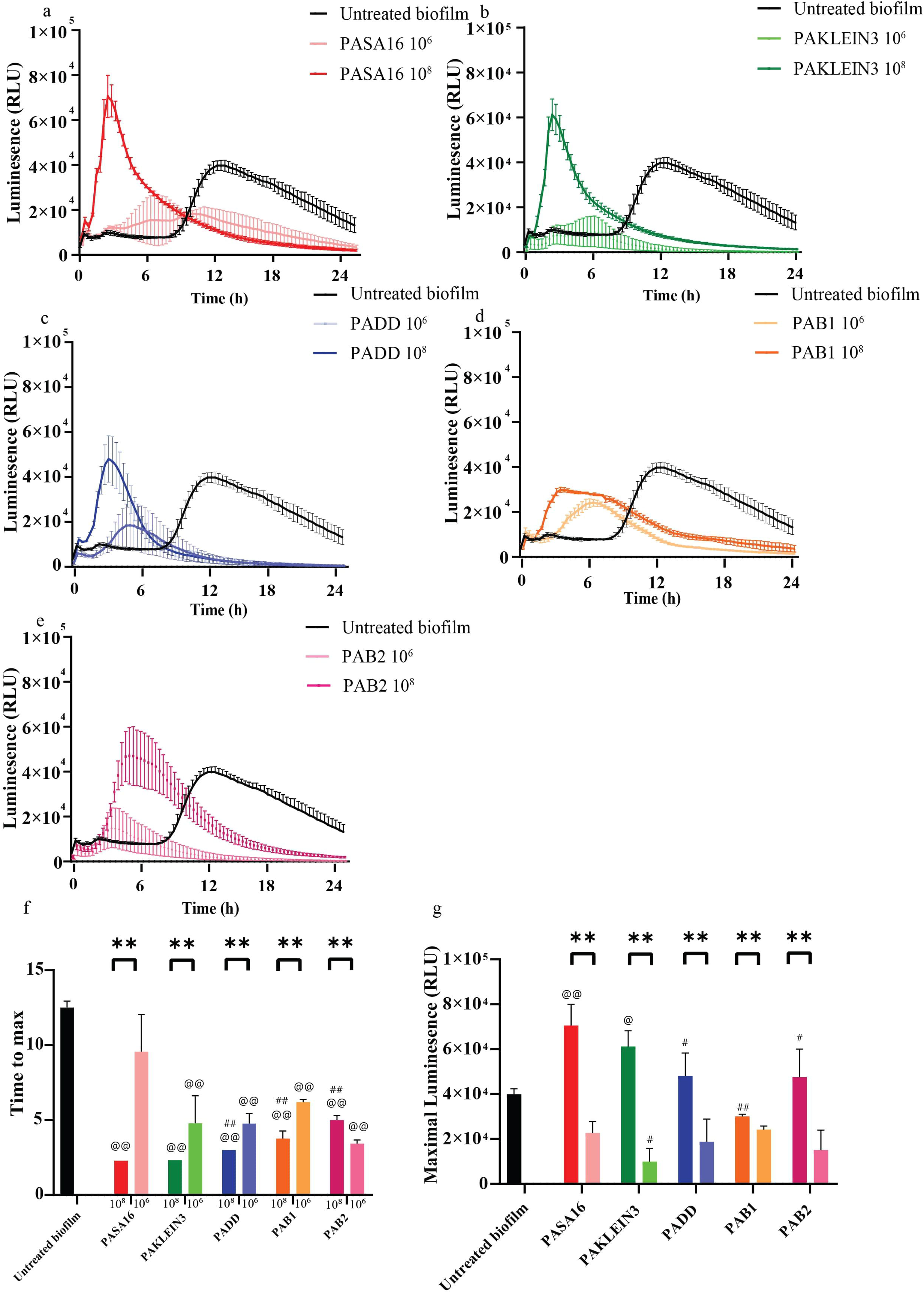
Assessing extracellular ATP change in phage-treated biofilms. Comparing *P. aeruginosa* biofilm extracellular ATP with five different phages at 10^8^ and 10^6^ PFU/ml. **a** PASA16 phage, **b** PAKLEIN3 phage, **c** PADD phage, **d** PAB1 phage, **e** PAB2 phage. **f** The time to max was calculated at different concentrations as described above. **g** the maximal luminescence was measured for each experiment. RealTime-Glo™ Extracellular ATP Assay was used. The results are the average of triplicates, presented as mean ± standard deviation. Representative results from two independent experiments. * *p <* 0.05, ** *p <* 0.001 by two-way Student’s t-test, comparing different titers of a given phage. # *p <* 0.05, ## *p <* 0.001 by one-way Student’s t-test, evaluating whether a given phage is significantly less effective than PASA16. @ *p <* 0.05, @@ *p <* 0.001 by one-way Student’s t-test, assessing whether each phage is significantly more effective than the control.

To better understand the assay’s dynamics, we examined lysed bacteria which demonstrated a dose response with a detection level of 10^5^ CFU/ml (Supplemental Fig. S3a). We then assessed planktonic bacteria. The addition of the reagent resulted in reduced OD_600_ measurements (Supplemental Fig. S3b) but was non-toxic to both bacteria and phages (Supplemental Fig. S1a), as per the manufacturer’s datasheet. When bacteria (initial CFU of at least 10^6^ CFU/ml) were exposed to phages, ATP was rapidly released into the medium. Control bacteria also eventually excreted ATP, but this occurred later than the phage-induced ATP release (Supplemental Figure 3c-h).

Upon evaluating three metabolic assays for detecting biofilm degradation, we found significant differences in their performance. The Extracellular Acidification Rate (ECAR) measurement demonstrated insufficient sensitivity to reliably detect changes in biofilm degradation. In contrast, both calorimetry and extracellular ATP (eATP) dye-based methods emerged as robust and sensitive alternatives (Table 2).

Calorimetry, while highly accurate, presents limitations due to the substantial cost of instrumentation and the requirement for anaerobic conditions during measurements. On the other hand, the eATP measurement technique showed high potential. Its high sensitivity, coupled with ease of use and adaptability to various experimental conditions, positions it as a promising cornerstone for future protocols in biofilm research and phage therapy studies.

### D. Genomic assays

#### 7. Extracellular DNA (exDNA) qPCR Assay

In our investigation of bacterial cell lysis, we employed quantitative PCR (qPCR) to measure extracellular DNA (exDNA) as a marker of phage-induced lysis. This approach focused on the housekeeping gene *rpoD* in *Pseudomonas aeruginosa* PA14 biofilms, comparing phage-exposed and unexposed samples over a 48-hour period. Our time-course analysis revealed that the most pronounced phage effect occurred at 24 hours post-treatment (Fig. 5a).

**Figure 5.**
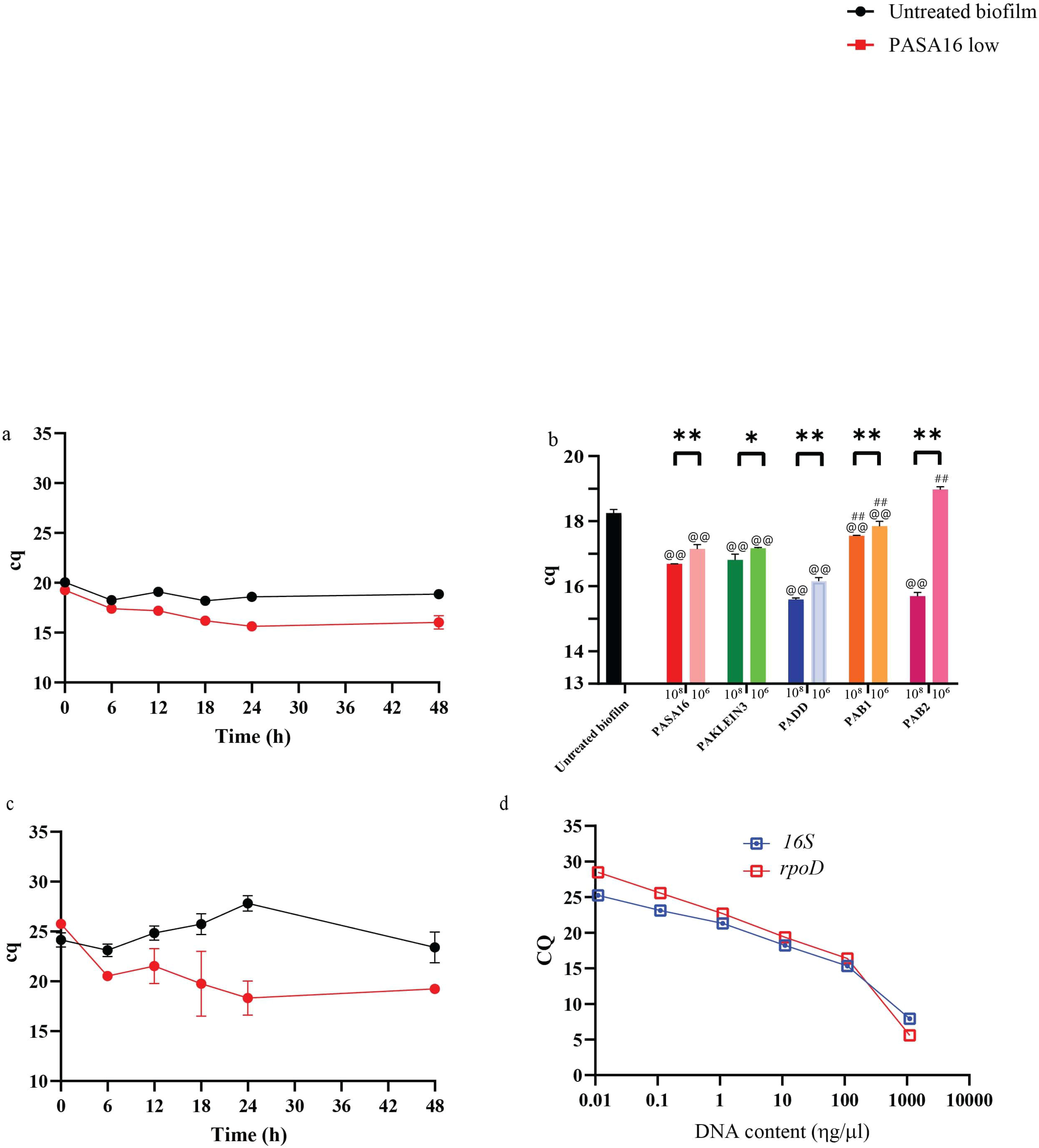
Assessing extracellular DNA change in phage-treated biofilms. This figure presents an assessment of extracellular DNA (exDNA) changes in biofilms treated with phages. Initially, a time-course analysis was conducted using the housekeeping gene *rpoD* as a marker to determine the optimal timing for qPCR analysis. PASA16 phage at a concentration of 10□ PFU/ml was used for this purpose. **b** Subsequently, biofilms were treated with five different phages at titers of 10□ and 10□ PFU/ml for 24 hours. Following treatment, qPCR was performed on filtered culture supernatants to quantify exDNA release by targeting the *rpoD* gene. **c** To further validate the generalizability of this method, a 48-hour kinetics study was conducted using *16S* rRNA gene primers, which confirmed that the most significant discrimination occurred at 24 hours post-treatment. **d** Additionally, calibration curves were constructed using exDNA isolated after antibiotic-induced lysis, utilizing primers for both rpoD and *16S* rRNA genes. The results are presented as mean ± standard deviation from triplicate measurements and are representative of two independent experiments. Statistical significance was determined as follows: \**p <* 0.05, \*\**p <* 0.001 by two-way Student’s t-test, comparing different titers of a given phage; #*p <* 0.05, ##*p <* 0.001 by one-way Student’s t-test, evaluating whether a given phage is significantly less effective than PASA16; @*p <* 0.05, @@*p <* 0.001 by one-way Student’s t-test, assessing whether each phage is significantly more effective than the control.

**Figure 6.**
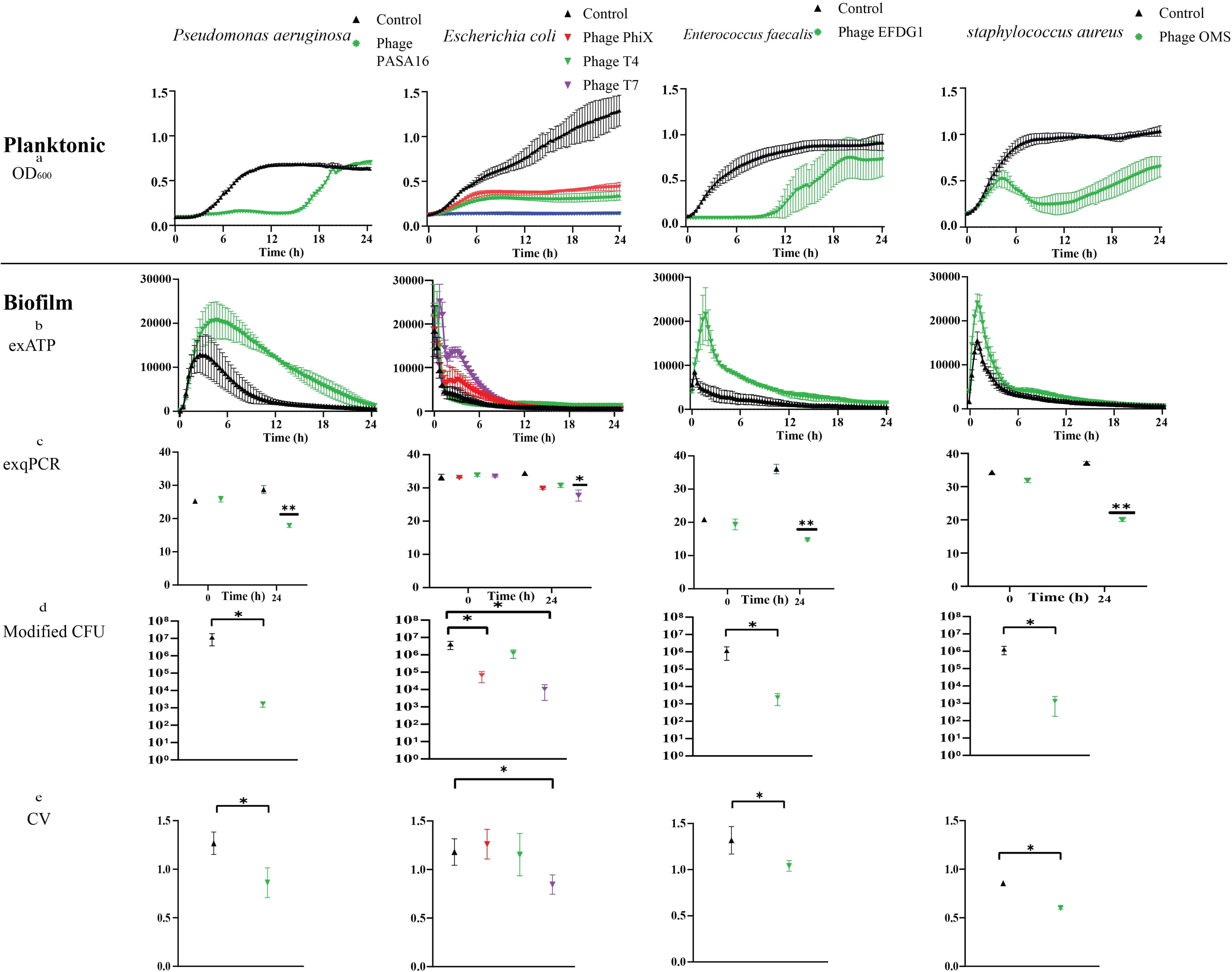
Testing a proposed protocol for biofilm-specific clinical phage microbiology on various bacteria. The protocol includes several methods for assessing biofilm formation and bacterial growth: eATP analysis, exDNA qPCR, Crystal violet staining, modified CFU analysis, and planktonic OD_600_ assessment. These methods were tested on both Gram-negative and Gram-positive bacteria, including *Pseudomonas aeruginosa* PA01 and *Escherichia coli DSM 13127* for Gram-negative, and *Staphylococcus aureus* R1 and *Enterococcus faecalis* (with the specific strain not provided) for Gram-positive, each with its respective phages at a titer of 10^8^ PFU/ml. The proposed protocol combines multiple techniques to provide a comprehensive assessment of biofilm formation and bacterial growth, offering insights into biofilm dispersal, matrix composition, biomass, viable bacterial counts, and planktonic growth. This multi-faceted approach allows for a more comprehensive evaluation of biofilm formation across different species, demonstrating the protocol’s versatility in studying clinically relevant pathogens known for forming biofilms.

Subsequent evaluation of various phage treatments demonstrated significant differences in quantification cycle (Cq) values. Untreated control groups exhibited Cq values approximately 90% higher than phage-treated groups (*p <* 0.001, Fig. 4b). All phage-treated groups showed reduced Cq values, with higher phage titers correlating to greater reductions (*p <* 0.05, Fig. 4b). These lower Cq values directly indicate higher exDNA levels, signifying more robust bacterial lysis.

To assess the generalizability of this method, we extended our analysis using 16S rRNA primers. Consistent with our *rpoD* findings, the kinetics of lysis using 16S primers also showed maximal effect at 24 hours (Fig. 5c). To standardize the relationship between Cq values and DNA quantity, we generated a calibration curve. This analysis revealed that DNA released through lysis was detectable at concentrations as low as 10□³ ηg/μL. A strong linear correlation between DNA content and qPCR detection was observed, with an average Pearson correlation coefficient of −0.79 and an R² value of 0.624 (Fig. 4d).

Our results demonstrate that exDNA quantification via qPCR can effectively distinguish between different phages and their respective titers. This method provides a sensitive and specific approach to quantifying phage-induced bacterial lysis. However, it is important to note its limitations: the process is labor-intensive and only allows for endpoint analysis rather than real-time monitoring (Table 2).

### E. Extracellular matrix assessment

#### 8. Immunofluorescent stains

To better understand extracellular matrix quantification, and to assess whether it is able to differentiate between different phage activities, EbbaBiolight 680 (https://www.ebbabiotech.com/products/ebbabiolight-680), a fluorescent dye, selectively binds to *P. aeruginosa* extracellular glucans, integral components of the biofilm extracellular matrix (ECM). The application of this dye followed the manufacturer’s instructions, with confirmation of its non-cytotoxic and non-phage-toxic nature (Supplemental Fig. S1a-b). Biofilm treated with each of the five phages at two concentrations underwent fluorescence measurement using a plate reader (Excitation at 540nm and absorbance at 680nm).

There was a 90% statistical reduction in AUC across all phage treatments in comparison to the control; notably, PAB2 exhibited an increase in absorbed fluorescence and is the only phage inferior to PASA16 (12.5%, *p <* 0.05, Supplemental Fig. S4 a-f). Further analysis revealed that only 60% of the five tested phages demonstrated a decrease in ECM at higher concentrations (*p <* 0.05, Supplemental Fig. S4 a-f).

#### 9. Scanning Electron Microscopy (SEM)

For the visualization of both biofilm viability and biomass, SEM was used in a magnitude of X5k, and X20k (Supplemental Fig. S5a). Phage was observed at a magnitude of X100k. Biofilm formed on a plastic surface was treated with the phage panel described above at two different concentrations. As there is no proved quantification method, it is qualitatively observed that out of all the tested phages, most phages displayed enhanced biofilm degradation at the higher phage titer. It can be assessed visally that the most effective phage was PASA16 as it has reduced biofilm presence in both titers used.

Although both methods appear promising, EbbaBiolight demonstrated limited sensitivity in our assay. While SEM is integral to the study of biofilm-phage interactions, it lacks quantitative precision and is too costly and labor-intensive for clinical biofilm phage therapy (Table 2).

### F. Other essays

#### Utilizing Bacterial Fluorescence

In this method, we utilized a Green Fluorescent Protein (GFP)-expressing strain of *P. aeruginosa* PA14 with a non-bacterial and non-phage toxic dye (Supplemental Fig. S1a-b). Aiming to mask the fluorescence of planktonic bacteria, using the Allura red dye. Therefore, enabling selective detection of green fluorescence emitted only by adherent bacteria forming the biofilm.

No statistically significant differences were found among the five applied phages (*p-value* not significant, Supplemental Fig. S6a-b).

To assess the method’s efficacy in measuring biofilm establishment, we conducted a proof-of-concept test with one phage, PASA16. While no differences were observed using the commonly employed OD_600_ nm for estimating bacterial concentration (*p-value* not significant, Supplemental Fig. S6c), fluorescence measurements revealed that PASA16-treated biofilm exhibited slower growth compared to the untreated control biofilm (*p-value* < 0.01, Supplemental Fig. S6d). another major limitation of this method is the need to have a GFP emitting bacteria.

#### Utilizing Bacterial Vibration

This system, developed by Resistell© (https://resistell.com/), attempts to quantify bacteria by measuring their vibration. While the method shows promise and is user-friendly, it requires labor-intensive calibration before each experiment. It has not yet produced reproducible results, primarily due to the limitation of having only one measurement chamber—a challenge that is expected to be addressed in future versions.

## Discussion

The findings from this study shed light on potential methodologies for assessing phage activity against biofilms, necessary due to the lack of activity of methods such as growth curve analysis, commonly used in planktonic bacteria(Yerushalmy et al., 2023).

Metabolic approaches for evaluating bacterial viability within biofilms have been previously established(Wadsö et al., 2017; Wilson et al., 2017). While enzymatic assessments have been applied in Biolog© systems(Cruz et al., 2021), isothermal calorimetry, and eATP assessment are yet to see wide clinical implementation, possibly due to initial high implementation costs or simply the lack of need. These techniques may find important applications in clinical settings, particularly in the context of phage therapy.

Other promising techniques, such as qPCR for exDNA, have been demonstrated to enumerate bacterial cell lysis in both planktonic(Hazan et al., 2016b) and biofilm-associated bacteria(Azeredo et al., 2017; Klein et al., 2012) Given the widespread use of qPCR during the Covid-19 pandemic, the device and technique should be within reach(Wang et al., 2020). One Significant limitation of this method is the requirement for filtration for all samples.

The conventional Colony Forming Unit (CFU) method, typically employed post-sonication, proved insufficient in our specific study of phage-infected *P. aeruginosa* biofilm, potentially due to the biofilm’s high viscosity and inadequate mechanical separation(Azeredo et al., 2017; Li et al., 2014) CFU, known for not accurately representing the fraction of detached live cells, encounters limitations in discerning viable but non-culturable (VBNC) cells(Azeredo et al., 2017; Li et al., 2014) To address the mechanical separation challenge, we implemented a subtle modification by cultivating the biofilm on stainless steel washers, which were then directly transferred to the sonication bath within sterile tubes. This may also be achieved using a 96-peg lid plate, which can next be moved to a different medium(Racenis et al., 2023) Additionally, in cases involving prosthetic treatment, the same material as the patient’s clinical infection site could be utilized for more targeted analysis. Crystal Violet (CV) emerged as a valuable alternative, offering sensitivity without being cost-prohibitive or labor-intensive. However, the fact that it measures biomass and the absence of a standardized protocol for implementation remain drawbacks(Azeredo et al., 2017; Stepanović et al., 2007).

We explored a metabolic approach using the Seahorse XFe96 Analyzer© to measure the Extracellular Acidification Rate (ECAR), initially designed for eukaryotic cell metabolism. ECAR correlates with glycolysis(S. Bhatia et al., 2021), while OCR (Oxygen Consumption Rate) reflects mitochondrial respiration(Caro-Maldonado et al., 2014). Though prior studies show *P. aeruginosa* biofilm metabolism is impacted by toxins(Mclamore et al., 2010), our attempts to measure phage activity on biofilms were inconclusive. The Seahorse XFe96 Analyzer may be relevant for phages inhibiting bacterial glycolysis, but its 12-hour limitation is a challenge for prolonged biofilm infection assessments. This is the first published trial of this method for this purpose.

To assess the extracellular matrix in phage-infected biofilms, we used Biolight 680, a dye that binds to cellulose and amyloid components in biofilms(Choong et al., 2021; Eckert et al., 2022). However, Biolight 680 demonstrated limited sensitivity in our study. Despite this, it remains user-friendly and cost-effective, requiring only a fluorescent plate reader and posing no toxicity to bacteria or phages, allowing real-time assessment.

The Live/Dead stain with Laser Confocal Scanning Microscopy is a well-established method for generating 3D images of biofilms and assessing biofilm structure with green SYTO-9 for live cells and red propidium iodide (PI) for dead cells(Azeredo et al., 2017; Khalifa et al., 2016; Neu & Lawrence, 2014; Rakov et al., 2021). However, we found this method less sensitive for phage-infected biofilms, likely due to phage-induced bacterial lysis, making PI ineffective. Additionally, the cytotoxicity of SYTO9 limits this method to endpoint analysis(Chiaraviglio & Kirby, 2014), reducing its clinical reliability.

Scanning Electron Microscopy (SEM) provides accurate results without the need for fluorescent dyes, but the sample preparation and visualization are expensive and labor-intensive (Azeredo et al., 2017; Kotra et al., 2000). Although quantification methods exist, they are not widely adopted(Vyas et al., 2016).

Bacterial viability was assessed using the Resistell system, which measures motion via a laser beam reflected from a cantilever with attached bacteria. However, the system’s limitation of handling only one sample at a time restricts technical repetitions and direct treatment comparisons. Additionally, biofilm growth on the cantilever can cause bending, potentially affecting accuracy. These issues are being addressed by the manufacturer and are expected to be resolved in the next device version. As a result, we have not yet obtained conclusive results on bacterial phage susceptibility within biofilms.

Notable approaches for future protocols include methods that can differentiate between 75-100% of inter-phage titers, distinct phages, and untreated controls, as well as phages inferior to PASA16. Promising techniques involve metabolic analyses, such as calorimetry, and the assessment of metabolic byproducts like eATP. Additionally, molecular methods like exPCR for extracellular DNA and a modified CFU approach using washers combined with Crystal Violet staining have shown potential. Although calorimetry was not pursued due to cost constraints, the other methods demonstrated initial efficacy in distinguishing between different phages tested in various bacterial species, including a different *Pseudomonas aeruginosa* strain, *Escherichia coli* (another Gram-negative rod), the Gram-positive bacteria *Enterococcus faecalis*, and *Staphylococcus aureus*.

This study’s limitations arise from the challenge of assessing the Multiplicity of Infection (MOI) in biofilm experiments due to uncertainty about bacterial exposure to phages. As a workaround, phage concentration, rather than MOI, was quantified. To gauge phage efficacy accurately, a lab reference strain and qualified phage titer were utilized, aiming to evaluate pure efficacy at similar concentrations, though acknowledging the need to consider different EOP between phages in clinical settings. Focusing on a single bacterial species and specific phages underscores their strain and phage specificity in reducing *Pseudomonas* biofilm biomass(Chang et al., 2019; Chegini et al., 2020) The study employed single prototype testing across all method categories, aiming for future validation with clinical strains, especially given antibiotic use in phage therapy. Despite efforts to standardize plastic or biofilm growth, inevitable variations occurred in well areas and plastic manufacturers. Establishing correlations between *in vitro* biofilm predictions and clinical outcomes is crucial for advancing the understanding of phage therapy efficacy.

In conclusion, the development of effective methods for matching phages to biofilm infections is crucial for advancing phage therapy. Future research should focus on refining and validating promising techniques to overcome current limitations. By leveraging these advancements, we can enhance our ability to tailor phage treatments to specific biofilm infections, ultimately improving therapeutic outcomes.

## Materials and Methods

All methods described herein were used in all relevant assays tested in this manuscript.

### Bacterial Strains and Culture Conditions

*Pseudomonas aeruginosa* strain PA14 was used as the primary model microorganism in this study, alongside *P. aeruginosa* PA14 GFP, *P. aeruginosa* PAO1, *Enterococcus faecalis*, *Staphylococcus aureus* R1, and *Escherichia coli DSM 13127*, all obtained from lab stock. These bacteria were routinely cultured in Luria-Bertani (LB, BD Franklin Lakes, New Jersey) broth or on LB agar plates (1.5% agar) at 37°C with shaking at 200 rpm for liquid cultures, or Brain Heart Infusion (BHI, BD) for *Enterococcus faecalis*. For biofilm formation, 180 µL of overnight grown bacteria were inoculated into each well of a 96-well plate, unless otherwise specified, and then incubated statically at 37°C for 24 hours to allow biofilm development (Müsken et al., 2010). Bacterial stocks were maintained at −80°C in LB broth containing 25% glycerol for long-term storage while working cultures on LB agar plates were stored at 4°C for up to one week.

### Determination of bacterial concentration

The determination of bacterial concentrations followed the standard Colony Forming Unit (CFU) assay method. Initially, the biofilm underwent washing with a saline solution and subsequent mechanical removal using a sterile tip. The contents of each well were then transferred into a 1.5 ml tube, following a previously established protocol (Shlezinger et al., 2019) The tubes were placed in a SONOREX sonication water bath (Bandelin, Berlin, Germany) for 5 minutes to effectively disrupt the biofilm. Subsequently, 5 µl of 10-fold serial dilutions were plated on LB agar plates. After 24 hours of incubation at 37□, colonies were counted, and the bacterial concentration was calculated in terms of CFU/ml.

For the modified washers CFU method, biofilm growth occurred on a Standard Flat stainless steel Washer (Fajoeda, Brazil) with these properties, M3 /0.118″; Outer Dia: 7mm/0.276″ Thickness: 0.5mm/0.02″, for 24 hours at 37□ in LB medium. Prior to phage application, the washer was transferred to a well with fresh LB medium. Preceding sonication, the washer underwent a saline wash and was then moved to a 15-ml tube. Subsequent dilutions were carried out from the saline solution, following the same procedure as described above.

### Phage used and determination of phage concentration

*P. aeruginosa* phages (Table 1) isolation is described in different publications(Alkalay-Oren et al., 2022; Rimon et al., 2025). Other phages used were *E. faecalis* targeting phage EFDG1(Khalifa et al., 2015), and *S. aurous* targeting phage OMS1(Porat et al., 2021) and *E. coli* targeting phages PhiX, T4, T7(Dunn et al., 1983; Romeyer Dherbey & Bertels, 2024; Wichman & Brown, 2010). Phages were stored in liquid LB at 4°C. The phage concentration was determined according to the PFU assay method. Phage suspensions have undergone 10-fold dilutions. 200 µl of overnight grown bacteria were put in 4 ml 0.5% agarose on LB agar plates to make bacterial lawns. 5 µl of each phage dilution was spotted twice on the bacterial lawns. The plates were dried and incubated aerobically for 24 hours. The number of plaques was counted, and the initial concentration of PFU/ml was calculated as described previously(Adams, 1959) If not specified otherwise, phages were grown to an initial concentration of 10^8^ PFU/ml.

### Growth curves

Phage activity on planktonic bacteria or biofilm was evaluated by inoculating logarithmic PA14 (at 10^7^ CFU/ml) or pre-grown biofilm with each unique phage sample. Turbidity, indicative of bacterial growth, was measured using a plate reader (Synergy; BioTek, Winooski, VT) at OD_600_ nm. The measurements were conducted at 37°C with 5-second shaking intervals every 20 minutes over a period of 24 hours.

### Calorimetry

For this purpose, calScreener**®** (Symcel, Solna, Sweden) was used, and biofilm was grown in designated activated Symcel-manufactured plates, as instructed by the manufacturer. 20 µl phages were added, and disposable plastic wells were placed in titanium chambers and sealed using a screwdriver. The heat measurement started after 1 hour due to the calScreener® initiation protocol. Data was analyzed using the supplied software.

### Extracellular acidification rate (ECAR)

For this purpose, the Seahorse XFe96 Analyzer (Agilent, Santa Clara, CA) was used with the company-supplied plates. Overnight culture of *P. aeruginosa* PA14 was grown in the Cell Plate (Agilent) for 24 hours. The utility plate was put in a calibrant medium as instructed by the manufacturer. Before treating the biofilm with phages, the biofilm was washed in 0.9% NaCl, and 180 µl of Running DMEM medium (Agilent) was put in the wells according to manufacturers’ instructions. 20 µl of either treatment was added in the different wells, and the plate was read by the machine every 12 minutes for 10 hours, the maximal period allowed by the manufacturer. Data was exported using Wave (Agilent) to PRISM (GraphPad, San Diego, CA).

### Extracellular ATP measurement

Biofilm cultures were established using standardized protocols. To each culture, 10 µl of either bacteriophage suspension or 0.9% NaCl (Saline) was added, bringing the final volume to 75 µl. The RealTime-Glo™ Extracellular ATP Assay (Promega, Madison, WI) was then applied according to the manufacturer’s instructions, with 25 µl of the assay reagent added to each sample(Ihssen et al., 2021). Luminescence measurements were recorded at 20-minute intervals using a Synergy plate reader. This assay allows for real-time monitoring of extracellular ATP levels, which can serve as an indicator of bacterial lysis and phage activity(Ihssen et al., 2021; Y. Liu et al., 2016). For calibration, *exATP* amounts bacterial lysis was achieved with carbenicillin at 50 µg/ml. Subsequently, lysate was serially diluted and assesed as described above.

### Extracellular DNA measurement

Biofilm was cultured following established procedures, and 20 µl of either phage or 0.9% NaCl was added. Quantification of relative extracellular DNA (*exDNA*) was performed on cell-free supernatants, filtered through a 0.22-μm pore size filter (Merck Millipore Ltd., Ireland). Real-time PCR was employed, utilizing specific primers for the *rpoD* (ACCGTCGTGGCTACAAATTC, GGCGATCTTCAGTACCTTGC(Hazan et al., 2016b)) and *16S* (CCTACGGGNGGCWGCAG, GACTACHVGGGTATCTAATCC(Klindworth et al., 2013)) genes, as previously described (Hazan et al., 2016b) The real-time PCR was conducted in a final volume of 25 µl, using the iQ™ SYBR® Green Supermix (Biorad, Hercules, CA) and the CFX96 Touch Real-Time PCR Detection System (Biorad). For calibration, *exDNA* amounts corresponding to nanodrop-measured DNA quantity, bacterial lysis was achieved with carbenicillin at 50 µg/ml. Subsequently, *exDNA* was filtered and quantified using nanodrop (Thermo Fisher).

### Biolight 680

(Ebba Biotech AB, Solna Sweden) was diluted to a stock solution according to the manufacturer’s instructions, Biolight 680 1:1000, and stored in the dark. Biofilm was grown on a 96-well plate and treated as described above. Biofilm was rinsed and 20 µl of phage treatment was applied as described, together with 20 µl of 10X Biolight 680. Fluorescence was measured by a Synergy plate reader (BioTek, Winooski, VT): 540 nm excitation and 680 nm emission for the Biolight 680 stain.

### Crystal Violet

Biofilm was grown on a 96-well plate and treated as described above. CV (Sigma Aldrich, St. Louis, MO) was used as previously described(Shteindel et al., 2019), 96 well plates were washed with 0.9% NaCl, then stained with 0.1% crystal violet solution (125 μl per well) incubated in ambient conditions for 15 min, and then emptied. Excess dye was rinsed twice in a water bath. The plate was dried in a 1.5 ml tube at 37 °C for 1 hour and extracted with 30% acetic acid (100 μl per well), and the extracted crystal violet was transferred into a new 96-well plate and its intensity was measured at OD_550_ nm.

### Live Dead stain

Biofilm was grown on a 96-well plate and treated as described above. The 96-well plate was washed and then stained using Live/Dead cell viability kits (Life Technologies, Waltham, MA) according to the manufacturer’s instructions. The fluorescence emissions of the samples were detected by using a Zeiss LSM 410 confocal laser microscope (Carl Zeiss). Red fluorescence was measured at 630 nm and green fluorescence was measured at 520 nm. Horizontal plane optical sections were made at 5-μm intervals from the surface outward and the images were displayed individually. The microscopy slices were combined into a 3D image using Bio-formats and UC± plugins (ImageJ 1.49G). The stained biofilms were examined using a confocal microscope and analyzed using ImageJ 1.49G software (http://imagej.nih.gov/ij/). %live of total bacteria was calculated as previously described(Robertson et al., 2019).

### Scanning electron microscopy (SEM)

Biofilm samples were prepared for SEM using Karnovsky’s fixative (2% PFA, 2.5% glutaraldehyde in 0.1 M cacodylate buffer, pH = 7.4). Samples in fixative were put for 4 h at room temperature, followed by 1:2 diluted Karnovsky’s fixative overnight at 4◦C. Then, the biofilm samples were post-fixed in 1% OSO4 in 0.1 M cacodylate buffer for 2 h and dehydrated in a graded series of alcohols, followed by Critical Point Drying (CPD). After sputter coating with Pd/Au, biofilm samples were observed by a SEM-trained technician (FEI, Quanta 200; CPD—Quorum Technologies, K850 Critical Point Drier; Sputtering—Quorum Technologies, SC7620 Sputter coater). Each well was treated and visualized in duplicate.

### Allure red

In this assay, we employed PA14 GFP as a fluorescent marker, while the concentration of Allure red dye (Sigma Aldrich) was meticulously calibrated following established procedures(Shteindel et al., 2019). Bacterial suspensions were prepared in distilled, deionized water (DDW) and then appropriately diluted to achieve an OD_600_ nm of 0.1. Subsequently, 100 µl of various Allure red dye concentrations were introduced into designated wells, and immediate bottom fluorescence readings were recorded using a plate reader set at an excitation wavelength of 395 nm and an emission wavelength of 509 nm. Post-calibration, a dye concentration of 0.8 mg/ml was deemed optimal, as it effectively reduced Relative Fluorescence Units (RFU) below 100 RFU, a criterion established in previous studies(Shteindel et al., 2019) (see Supplemental Fig. S6a). This validated dye concentration of 0.8 mg/ml was consistently applied in subsequent experiments, and RFU measurements were taken accordingly.

### Cell vibration

For cell vibration experiments, Resistell AG (Muttenz, Switzerland) provided the necessary equipment. Initially, the chamber was filled with filtered LB, and the pre-activated cantilever was carefully adjusted as per the manufacturer’s instructions. Subsequently, a baseline measurement was conducted over a duration of 10 minutes. Following this, bacteria were affixed to the cantilever, and measurements were taken over a 30-minute period. After this initial phase, either phage or a control substance was introduced, and the vibration was continuously monitored for a total of 20 hours. The acquired data were then subjected to analysis using Resistell’s dedicated software.

### Assessing dye’s bacterial toxicity and phage toxicity

In this assay, the bacterium used was *Pseudomonas aeruginosa* PA14, and the phage used was PASA16. Phage or bacteria were incubated overnight with the tested dyes or 0.9% NaCl solution, followed by CFU assay for bacteria and PFU assay for phages.

### Statistical analysis

Prism (GraphPad, version 8.0.2 (263), San Diego, CA) was used for the statistical analysis, graph formation, and design. The results were analyzed as the mean ± standard deviation in each experimental group and were repeated in two independent experiments. Statistical significance was calculated by a Student’s t-test with one or two-tailed unpaired *p-value*s (significance level: *p-value*s < 0.05).

## Data availability

The authors confirm that the data supporting the findings of this study are available within the article and its supplementary materials.

## Acknowledgment

We express our sincere appreciation to the Core Research Facility of the Hebrew University, Ein Karem Campus, and extend our gratitude to Dr. Eduard Berenshtein for facilitating access to TEM and SEM microscopy resources. Special thanks to Dr. Abed Nasereddin and Dr. Idit Shiff for their invaluable contributions to the deep sequencing procedures. We would also like to acknowledge the generous support of Symcel and Resistell, start-ups that graciously provided their products for a limited period without charge, enhancing the scope and depth of our research.

## Author contributions

All authors made significant contributions to the preparation of this manuscript.

AR, RNP, and RH conceptualized and drafted the manuscript, overseeing the funding acquisition process. RB and SCG have edited the manuscript. OY, HO, NK, LY, and SAO were instrumental in executing various methodologies. Supervision throughout the project was provided by YG and RH.

## Author Disclosure Statement

No competing financial interests exist.

During the preparation of this work, the authors used OpenAI-ChatGPT4 (https://chatgpt.com) and Grammarly (https://app.grammarly.com) to correct language and grammar. After using this tool/service, the authors reviewed and edited the content as needed and took full responsibility for the content of the published article.

