## Supplemental data for "Towards Clinical Phage Microbiology for Biofilm Infections: A Comparative Study and a Suggested Protocol"

**Supplementary materials:**

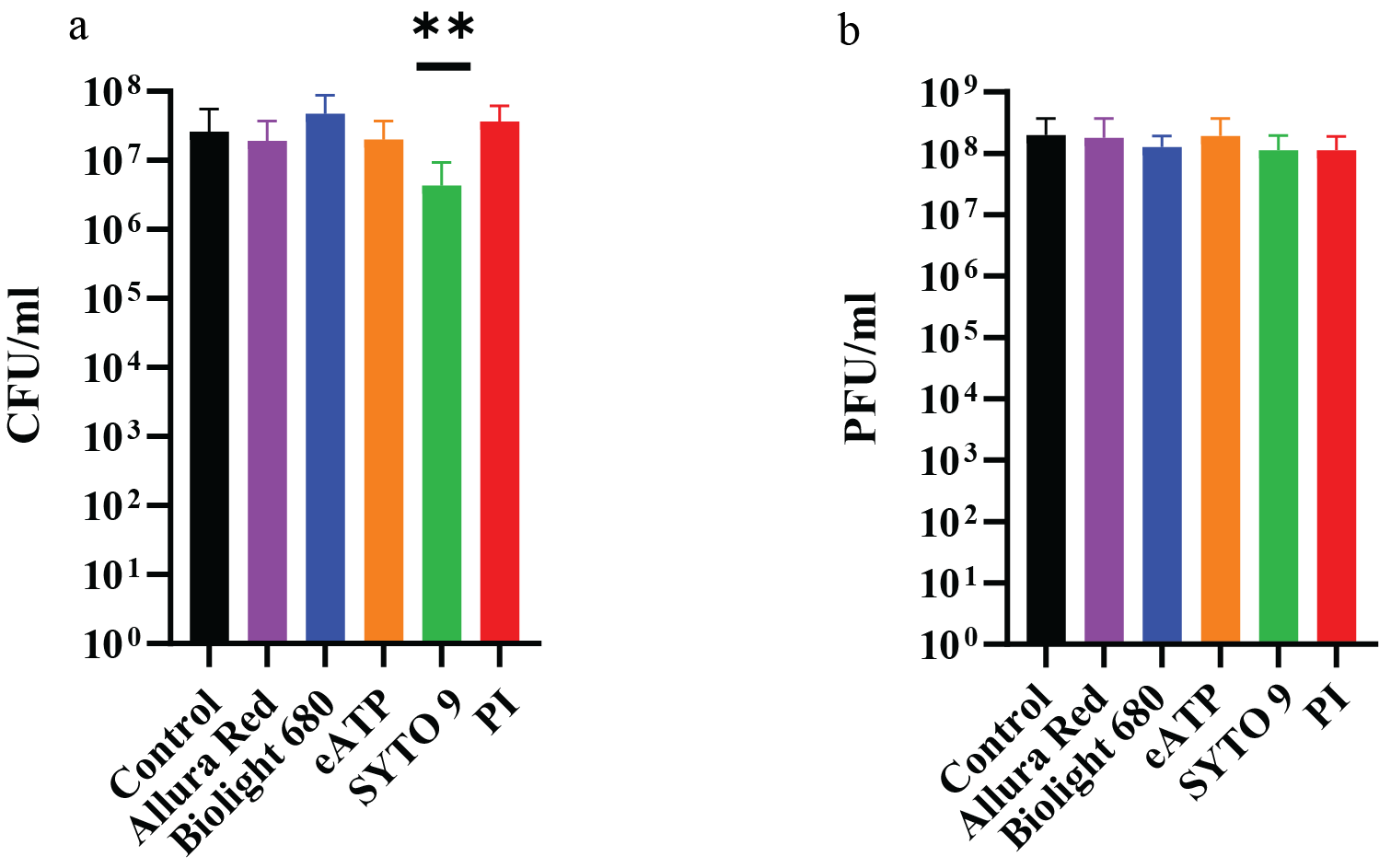

**Supplemental Figure S1 Bacterial and phage toxicity.** The assessment of bacterial and phage toxicity involved the utilization of various materials within the experimental tests, employing the bacterium *Pseudomonas aeruginosa* PA14 and the phage PAS16. In this evaluation, either the phage or bacteria were subjected to overnight incubation with the tested material or a 0.9% NaCl solution. Subsequent to the incubation period, CFU assays were conducted for the bacteria. **b** PFU assays were performed for the phages. Propidium Iodide (PI) was found to reduce bacterial viability. This comprehensive approach allowed for a thorough examination of the potential impacts of the materials on both bacterial and phage viability, providing valuable insights into their respective toxicities. The results are the average of triplicates, presented as mean ± standard deviation. Representative results from two independent experiments. ***p-value<*0.001 by 2-way Student t-test.

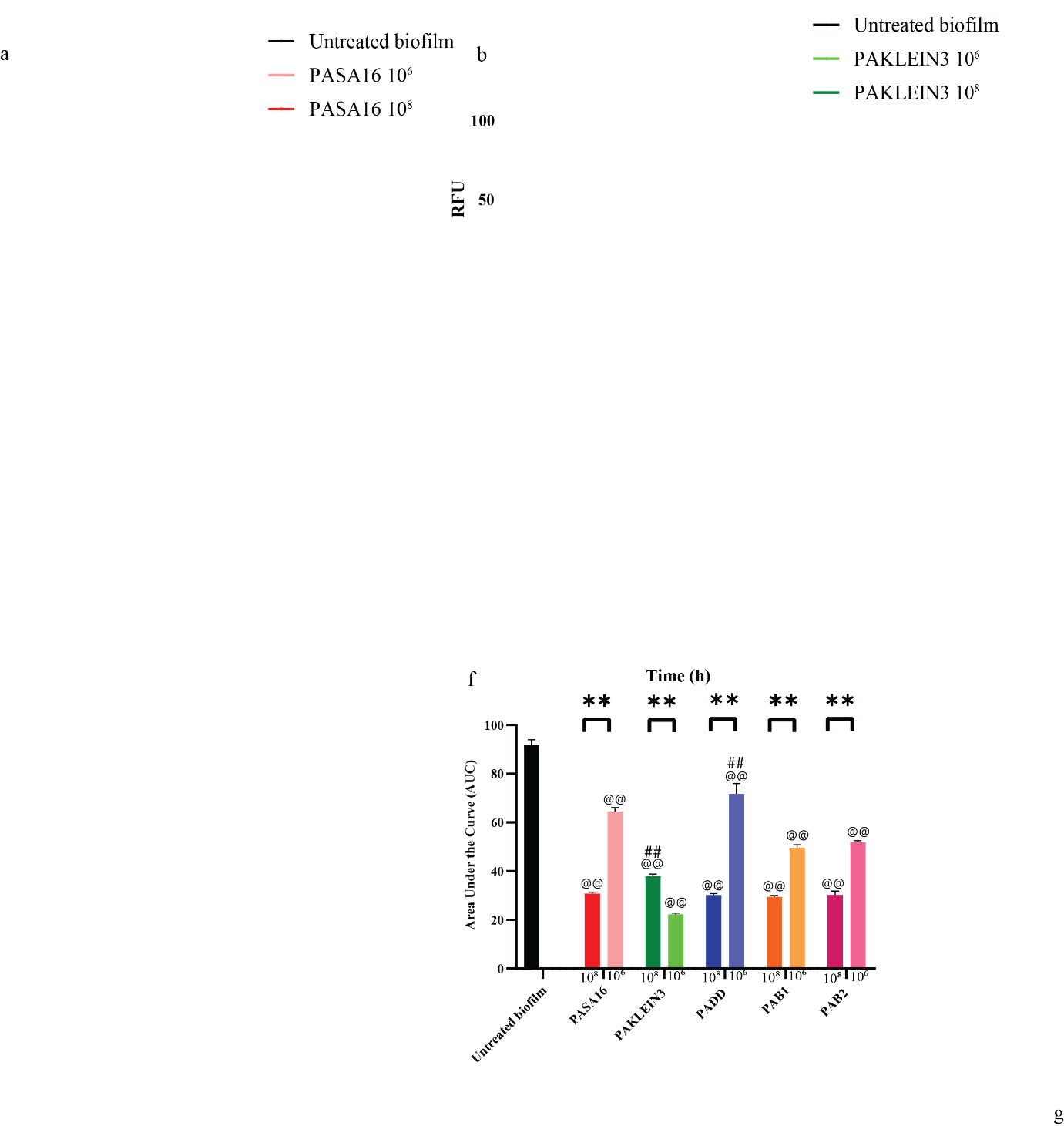

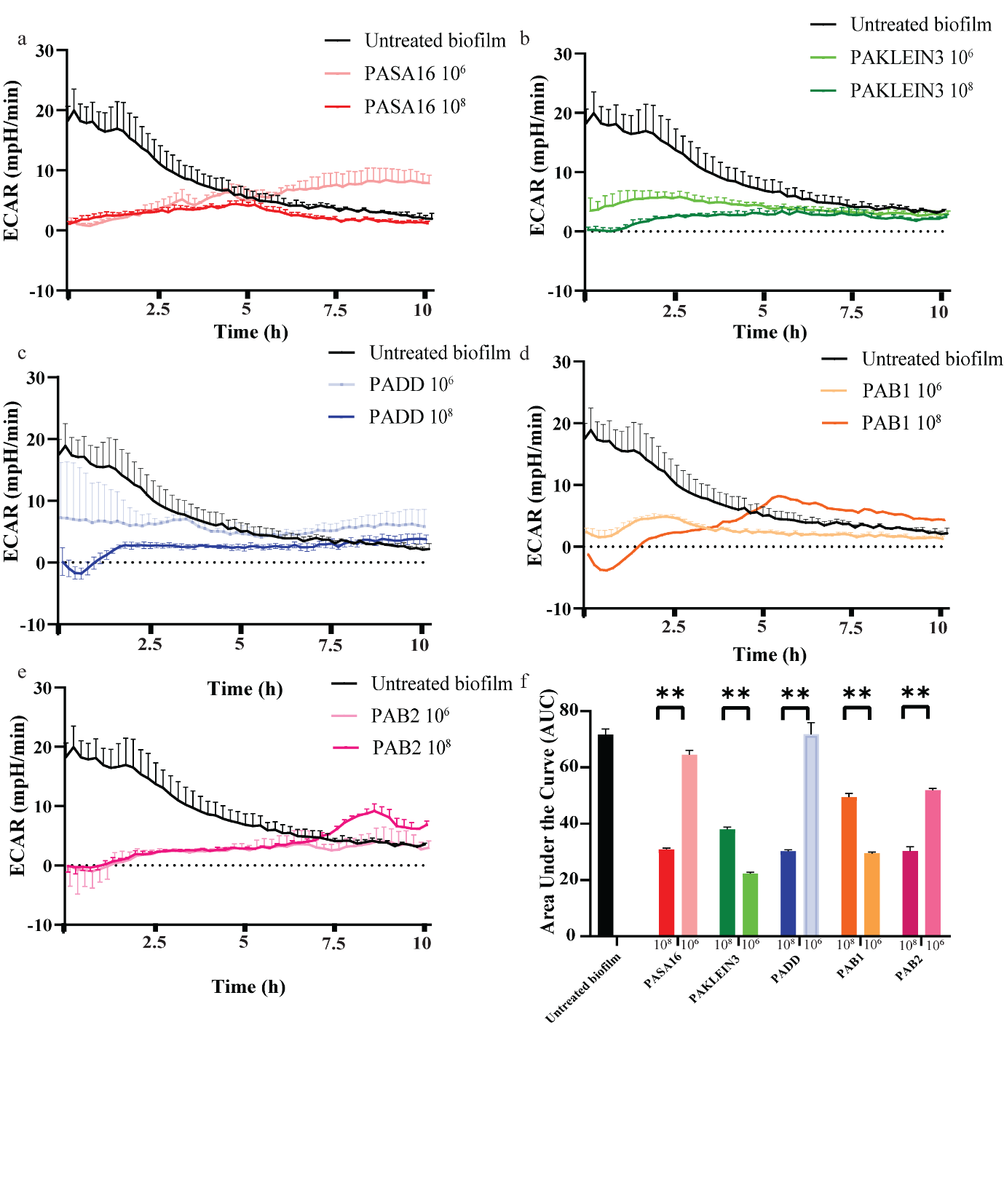

**Supplemental Figure S2. Extracellular acidification rate (ECAR) based phage biofilm effect comparison**. **a-e** Comparing H+ flux of a *P. aeruginosa* biofilm, treated with five different phages at a PFU of 10^8^ and 10^6^ . **a** PASA16 phage, **b** PAKLEIN3 phage, **c** PADD phage, **d** PAB1 phage, **e** PAB2 phage. ECAR was measured every 12 minutes, for 10 hours. **f** The AUC of phage activity was calculated at different concentrations (as described above). Notably, a consistent pattern of lower metabolic activity was observed in three out of five phages when a higher phage titer was applied. ECAR was measured using Seahorse XFe96 Analyzer^®^. The results are the average of triplicates, presented as mean ± standard deviation. Representative results from two independent experiments. * *p <* 0.05, ** *p <* 0.001 by two-way Student's t-test, comparing different titers of a given phage. # *p <* 0.05, ## *p <* 0.001 by one-way Student's t-test, evaluating whether a given phage is significantly less effective than PASA16. @ *p <* 0.05, @@ *p <* 0.001 by one-way Student's t-test, assessing whether each phage is significantly more effective than the control.

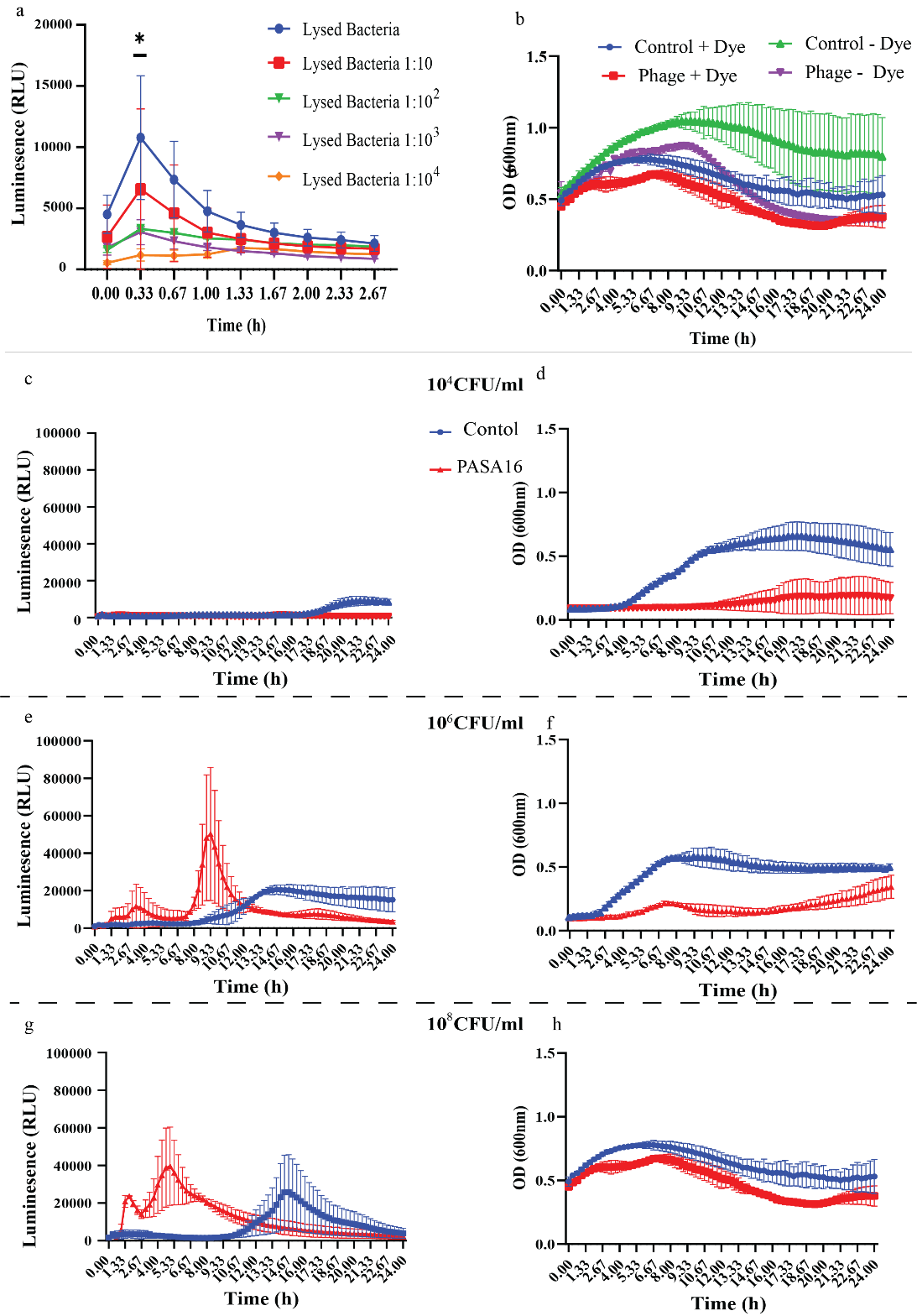

**Supplemental Figure S3. Extracellular ATP calibration. a** Heat-lysed bacteria were measured at various dilutions, demonstrating a dose-dependent response within the first hour of measurement. **b** Bacterial samples, both with and without phage, were assessed using OD_600_ measurements in conjunction with the RealTime-Glo™ Extracellular ATP Assay. While the addition of the assay resulted in reduced OD_600_ readings, it did not affect the CFU count (as shown in Supplemental Figure 1). **c-h** Multiple bacterial concentrations were tested, both in the presence and absence of phage, with simultaneous measurements of luminescence and OD_600_. The assay exhibited a detection threshold of 10^6^ CFU/ml. Notably, higher phage concentrations correlated with a shorter time to reach maximum signal intensity. The results are the average of triplicates, presented as mean ± standard deviation. Representative results from two independent experiments. **p-value<*0.05 by 2-way Student t-test.

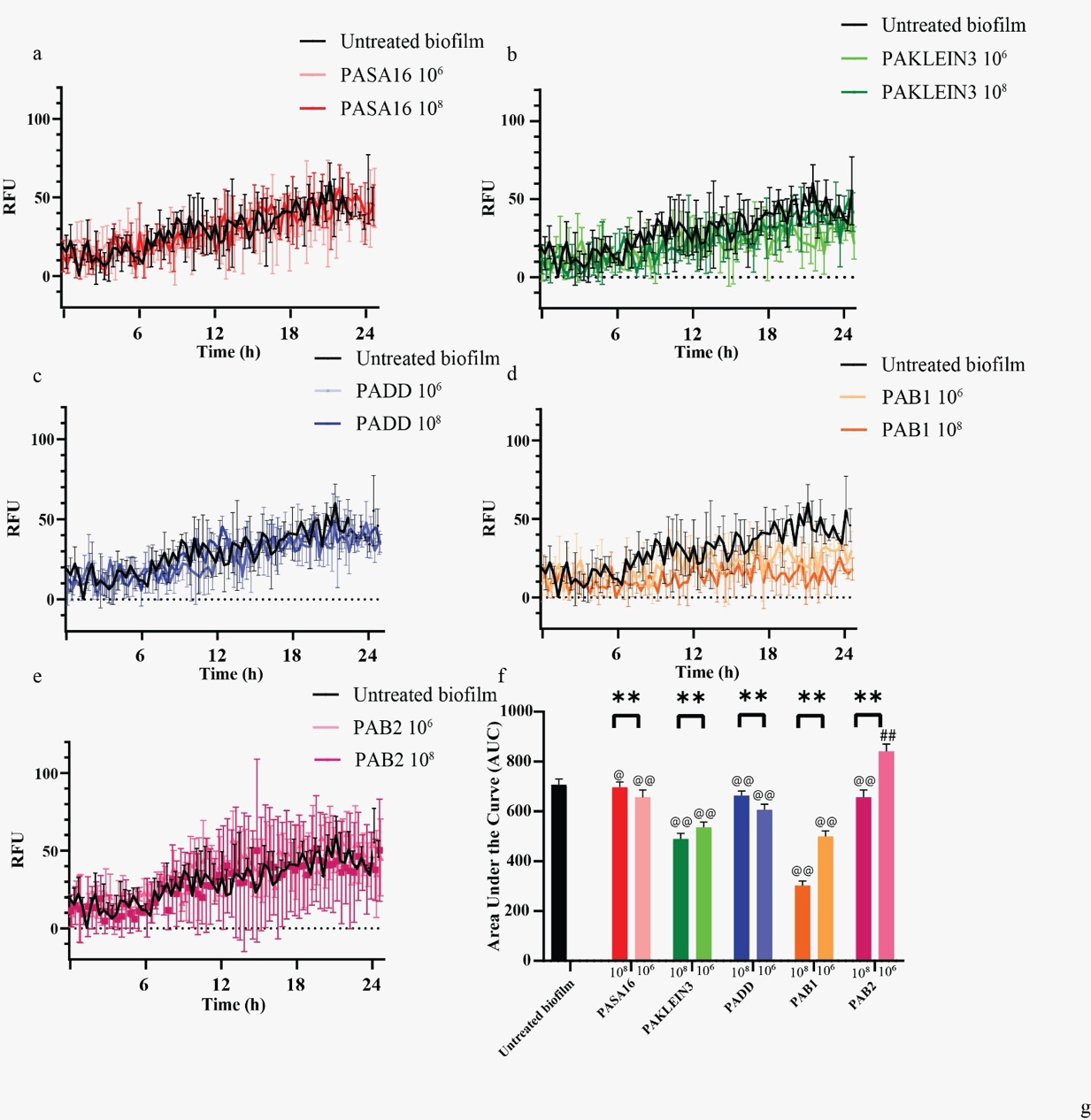

**Supplemental Figure S4. Monitoring extracellular glucan matrix to assess phage biofilm effect comparison. a-e** The evaluation of phage impact on the extracellular glucan matrix in PA14 biofilm was conducted through Biolight 680 fluorescent dye, comparing biofilms treated with five different phages at concentrations of 10^8^ and 10^6^ PFU/ml. The specific phages examined were: **(a)** PASA16 phage, **(b)** PAKLEIN3 phage, **(c)** PADD phage, **(d)** PAB1 phage, and **(e)** PAB2 phage. **(f)** The Area Under the Curve (AUC) was calculated to quantify phage activity at different concentrations. The resulting data demonstrated varied responses across the phages, with some exhibiting lower extracellular activity when exposed to a higher phage titer. Intriguingly, certain phages either showed no detectable activity or unexpectedly led to an increase in fluorescence. The results are the average of triplicates, presented as mean ± standard deviation. Representative results from two independent experiments. ** *p <* 0.001 by two-way Student's t-test, comparing different titers of a given phage. ## *p <* 0.001 by one-way Student's t-test, evaluating whether a given phage is significantly less effective than PASA16. @ *p <* 0.05, @@ *p <* 0.001 by one-way Student's t-test, assessing whether each phage is significantly more effective than the control.

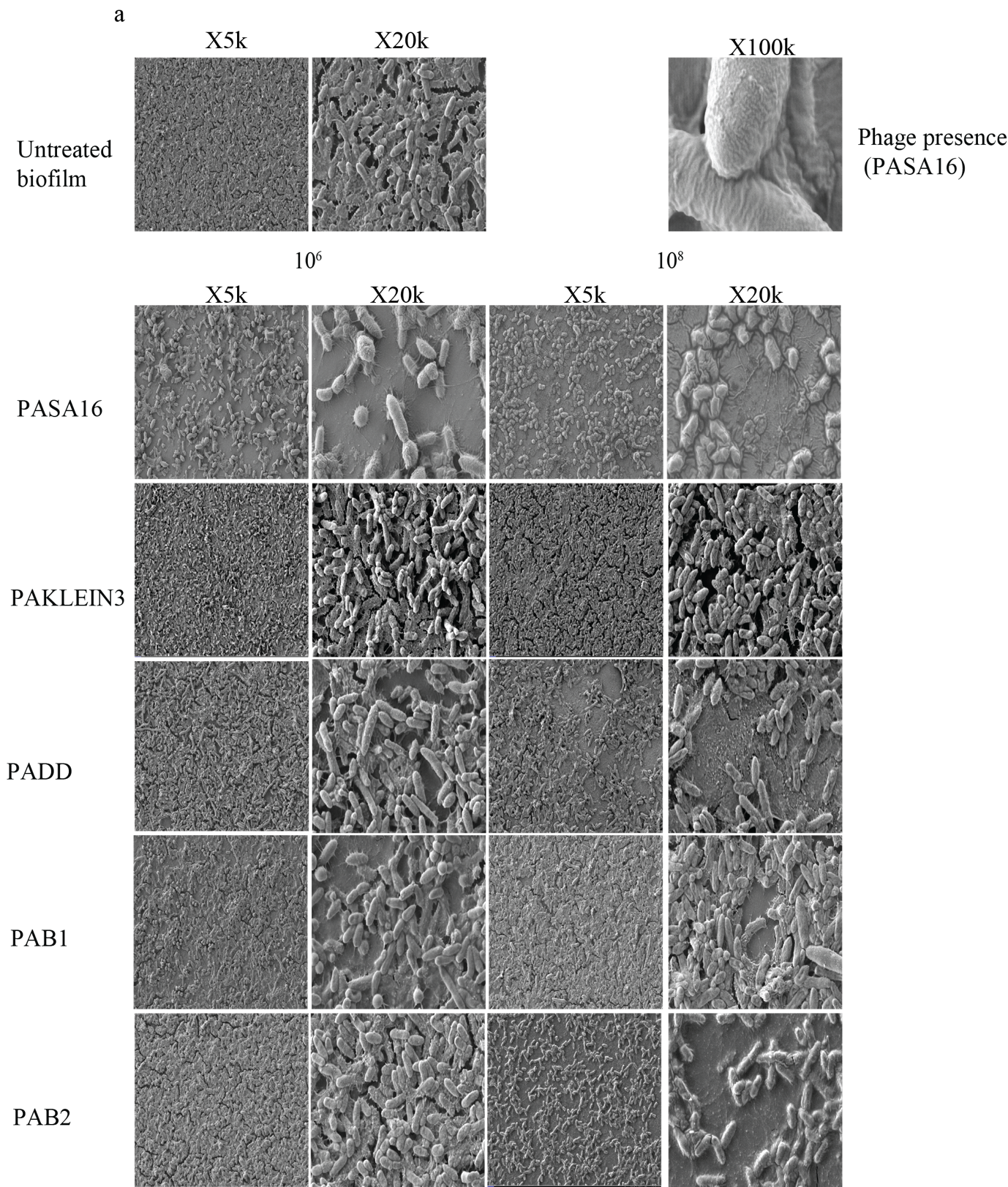

**Supplemental Figure S5. Assessment of Pre-grown Biofilm through Scanning Electron Microscopy (SEM) and Model Establishment a** Pre-grown biofilms were cultivated on plastic chambers and treated with five phages at varying concentrations. Samples were visualized using SEM at 5,000x and 20,000x magnifications The images shown are representative of two independent experiments.

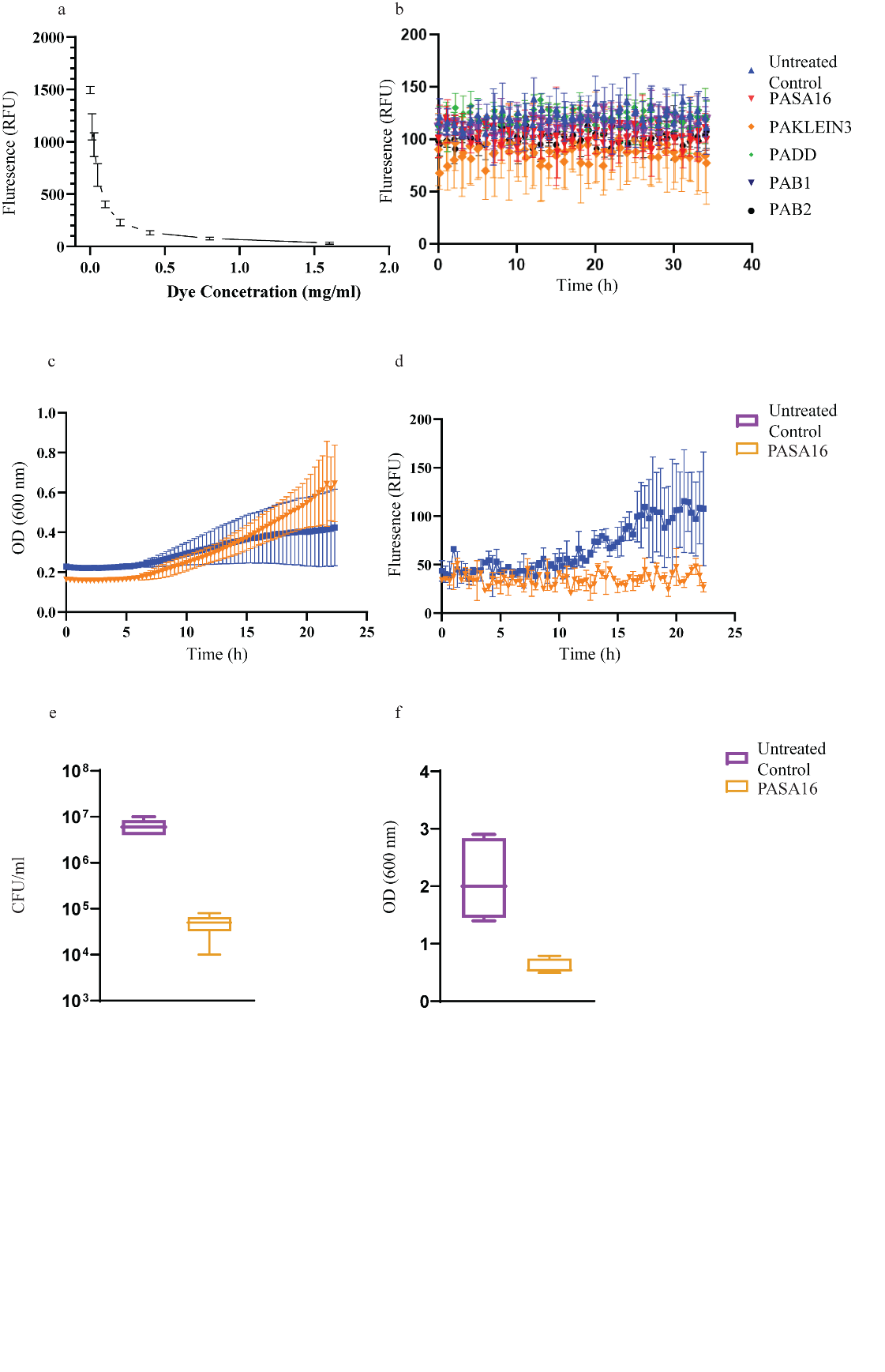

**Supplemental Figure S6. Allura red stain.** Emitted fluorescence, measured from *P. aeruginosa* PA14 GFP grown biofilm. By using Allura Red dye to mask the emission by the planktonic bacteria in the well, only the fluorescence emitted from surface-attached bacteria was absorbed. **(a)** Allura Red 0.8 mg/ml was used as it was the lowest dye concentration to achieve a sub-100 RFU in our calibration test. **(b)** PASA16, PAKLEIN3, PADD, PAB1, and PAB2 phage comparison, all at a concentration of 10^8^ PFU/ml, applied on a pre-grown biofilm. Application of PASA16 before the establishment of a biofilm: **(c)** RFU measured throughout biofilm establishment in the presence of Allura Red in the medium at a concentration of 0.8 mg/ml, **(d)** OD_600_ nm of bacterial growth throughout the process, in the presence of Allura Red at a concentration of 0.8 mg/ml. The results are the average of triplicates, presented as mean ± standard deviation. Representative results from two independent experiments.
